# Myelin-Weighted Core Network Connectivity Supports Trait-Level Future Thinking

**DOI:** 10.64898/2026.09.29.754335

**Authors:** Lucie S. Reed, Eirini Messaritaki, Craig Hedge, Carl J. Hodgetts, Carolyn B. McNabb, Derek K. Jones, Andrew D. Lawrence

**Affiliations:** School of Psychology, University of Birmingham, Edgbaston, Birmingham, B15 2TT; Cardiff University Brain Research Imaging Centre (CUBRIC), School of Psychology, Cardiff University, Cardiff, CF24 4HQ, Wales, UK; School of Psychology, Aston University, Birmingham, United Kingdom; Department of Psychology, Royal Holloway University of London, Egham, Surrey, TW20 0EX, UK; Department of Psychology, University of Edinburgh, 7 George Square, Edinburgh EH8 9JZ, UK

**Keywords:** Autobiographical memory, Default mode network, Episodic future thinking, Survey of Autobiographical Memory, Structural connectivity, Myelin

## Abstract

Why are some individuals able to recollect past events and imagine future events more richly than others? Here we examine how structural connectivity within a ‘core network’ supporting mnemonic cognition, substantially overlapping with the canonical default network, underpins inter-individual variation in autobiographical thought. We analysed state-of-the-art magnetic resonance imaging data from 109 adults collected on a high-gradient 3T Connectom scanner, in relation to their Survey of Autobiographical Memory (SAM) scores - a psychometric measure of self-reported episodic autobiographical, semantic, spatial, and future thinking abilities. We anatomically defined a core network, employed streamline tractography to derive participant-specific structural brain networks, and weighted their connections with the myelin water fraction derived from quantitative multicomponent relaxometry. We investigated associations between graph theoretical metrics of the core network and self-reported mnemonic abilities. We found a negative association between future thinking abilities and characteristic path length, and a positive association between future thinking abilities and mean clustering coefficient. Further analysis showed that the node-specific clustering coefficient of most core network nodes was positively correlated with future thinking scores, with frontotemporal nodes showing the strongest associations. Our findings indicate that both efficient integration and local segregation of the core network are important for trait-level future thinking.

**Author Summary:** People vary considerably in how richly they imagine autobiographical events. Although functional brain activity has been linked to individual differences in autobiographical cognition, less is known about how the structural organisation of this network contributes to these differences. Using advanced MRI data from 109 healthy adults, we examined whether the organisation of white matter connections within a core autobiographical memory network relates to self-reported memory and future thinking abilities. We found that individuals reporting stronger future thinking abilities exhibited shorter characteristic path length and higher clustering within the core network linked to autobiographical cognition. This suggests that individual differences in imagining the future may emerge from how this core network is structurally organised, indicating that both efficient integration and local segregation support trait-level future thinking.

## 1. Introduction

The human brain can be likened to a time machine, that allows events from one’s past and possible future to be experienced in the present (Tulving, 2001). This ability, known as autobiographical memory, varies substantially between individuals (Palombo et al., 2018). There is significant evidence for both highly superior (HSAM; LePort et al., 2012) and severely deficient (SDAM; Palombo et al., 2015) autobiographical memory, but more nuanced trait-level differences also exist between individuals (Berntsen et al., 2019; Palombo et al., 2013; Rubin, 2021). These differences are often marked by variation in functional activity and coactivation in a set of distributed neural regions known as the ‘core network’ that supports autobiographical functioning (Benoit & Schacter, 2015; Palombo et al., 2015; Rugg & Vilberg, 2013; Talbot et al., 2025). This network supports constructive mnemonic cognition, including recollection of past experiences, integration of semantic knowledge, construction of spatially coherent scenes, and simulation of plausible future events. It enables the flexible retrieval and recombination of episodic information which underpins autobiographical remembering, future thinking and other forms of mnemonic simulation (Andrews-Hanna et al., 2014; Benoit & Schacter, 2015; Hassabis & Maguire, 2007; Shapira-Lichter et al., 2013; Spreng et al., 2009; Vatansever, 2026).

The Survey of Autobiographical Memory (SAM; Palombo et al., 2013) captures stable individual differences in mnemonic traits. Distinct from performance-based measures and phenomenological ratings which assess characteristics of autobiographical events, the SAM measures everyday mnemonic abilities and stable individual differences that are difficult to capture in the laboratory. The questionnaire is comprised of four dimensions: episodic memory (the ability to remember personally experienced events), semantic memory (our general knowledge about the world), spatial memory (the awareness of previously encoded allocentric spatial contexts) and future thinking (the ability to imagine plausible personal future events).

Examining brain connectivity is crucial for understanding how the core network supports trait mnemonics, and non-invasive magnetic resonance imaging (MRI) is well suited for that goal. Yet few studies have investigated functional connectivity underpinning the SAM subscales. Sheldon et al. (2016) revealed that SAM-episodic scores were most strongly correlated with parahippocampal connectivity to posterior occipital and parietal cortices whereas semantic scores were most strongly correlated with parahippocampal connectivity to inferior and middle prefrontal regions. Teghil et al. (2022) also found significant correlations between SAM-episodic scores and BOLD signal change in the right hippocampus, right angular gyrus, posterior cingulate cortex and posterior cerebellum, whereas semantic scores were only associated with the right hippocampus. These studies provide some insights into the functional connectivity underpinning mnemonic traits; linking the different SAM dimensions to distinct connectivity patterns.

Brain function is shaped by structural connectivity (Jung et al., 2017; Mesulam, 1990; Suárez et al., 2020). MRI can quantify brain structural attributes that support brain function and may help explain individual differences in mnemonic abilities. White matter tracts support functional networks (Jung et al., 2017) and exhibit stable inter-individual differences (Owen et al., 2013; Karahan et al., 2022). Connectivity between pairs of brain regions involved in memory (Alexander-Bloch et al., 2013; Clark et al., 2022; Hodgetts et al., 2017; Williams et al., 2020) does not capture complex interactions between multiple brain areas. Treating the brain as a network of interconnected function-specific areas allows us to quantify how ensembles of brain areas work together to support cognition (Bullmore & Sporns, 2009; Sporns, 2010). Graph theory characterises large-scale network organisation and reveals possible communication mechanisms (Petersen & Sporns, 2015). Some studies have used graph theory methods to determine how *functional* networks relate to episodic memory (Geib et al., 2017a; Geib et al., 2017b; Ray et al., 2020; Schedlbauer et al., 2014). Only one study (Matyi & Spielberg, 2022) has examined structural connectivity in relation to mnemonic abilities (Penn Word Memory Test, Gur et al., 2010; Picture Sequence Memory Test, Dikmen et al., 2014). Performance on these tests correlated with node strength (i.e., the number of connections a brain region has within the network) of the medial temporal lobe and retrosplenial cortex. However, this study looked at a single measure of structural organisation and employed performance-based episodic memory tasks, likely capturing one aspect of remembering rather than the multi-faceted nature of autobiographical memory (Berntsen et al., 2019; McDermott et al., 2009; Roediger & McDermott, 2013).

We aim to understand the relationship between the 4 subscales of the SAM (episodic, semantic, spatial, and future thinking) and the organisational properties of the core network (Benoit & Schacter, 2015; Hassabis & Maguire, 2007) in healthy adults. We use state-of-the-art MRI data collected on a high-gradient 3T Connectom MRI scanner to reconstruct white matter pathways and generate structural subnetworks, with cortical regions as nodes and white matter tracts as connections (edges). Recognising the significance of myelin for brain function and efficient communication (Bells et al., 2017; Messaritaki et al., 2021; Nelson et al., 2026; Suminaite et al., 2019; Yeh et al., 2021), we use myelin measurements to assign connection strength. We calculate graph theoretical metrics that measure network integration and segregation. Metrics of integration index how effectively information is exchanged across regions within the network whereas metrics of segregation capture how specialised subnetworks support local processing (Mišić et al., 2014; Petersen & Sporns, 2015; Sporns, 2013). We examine the characteristic path length, global efficiency, mean clustering coefficient and mean nodal strength of the network-of-interest (Bullmore & Sporns, 2009). To derive network-wide measures for the latter two, their mean values over all the nodes of the network are used.

No previous study has examined how myelin-weighted structural network organisation relates to individual differences across the four dimensions of the SAM. We hypothesise that the SAM scores will be positively correlated with the mean clustering coefficient, mean nodal strength and global efficiency, and negatively correlated with the characteristic path length of the core network.

## 2. Methods

A graphical representation of the pre-processing and analysis pipeline is shown in Fig. 1. Multi-modal imaging and self-report data were collected as part of the Welsh Advanced Neuroimaging Database (WAND) study (McNabb et al., 2025) at the Cardiff University Brain Research Imaging Centre (CUBRIC). The data are publicly available and have been described in detail in the data release publication (McNabb et al., 2025).

**Figure 1:**
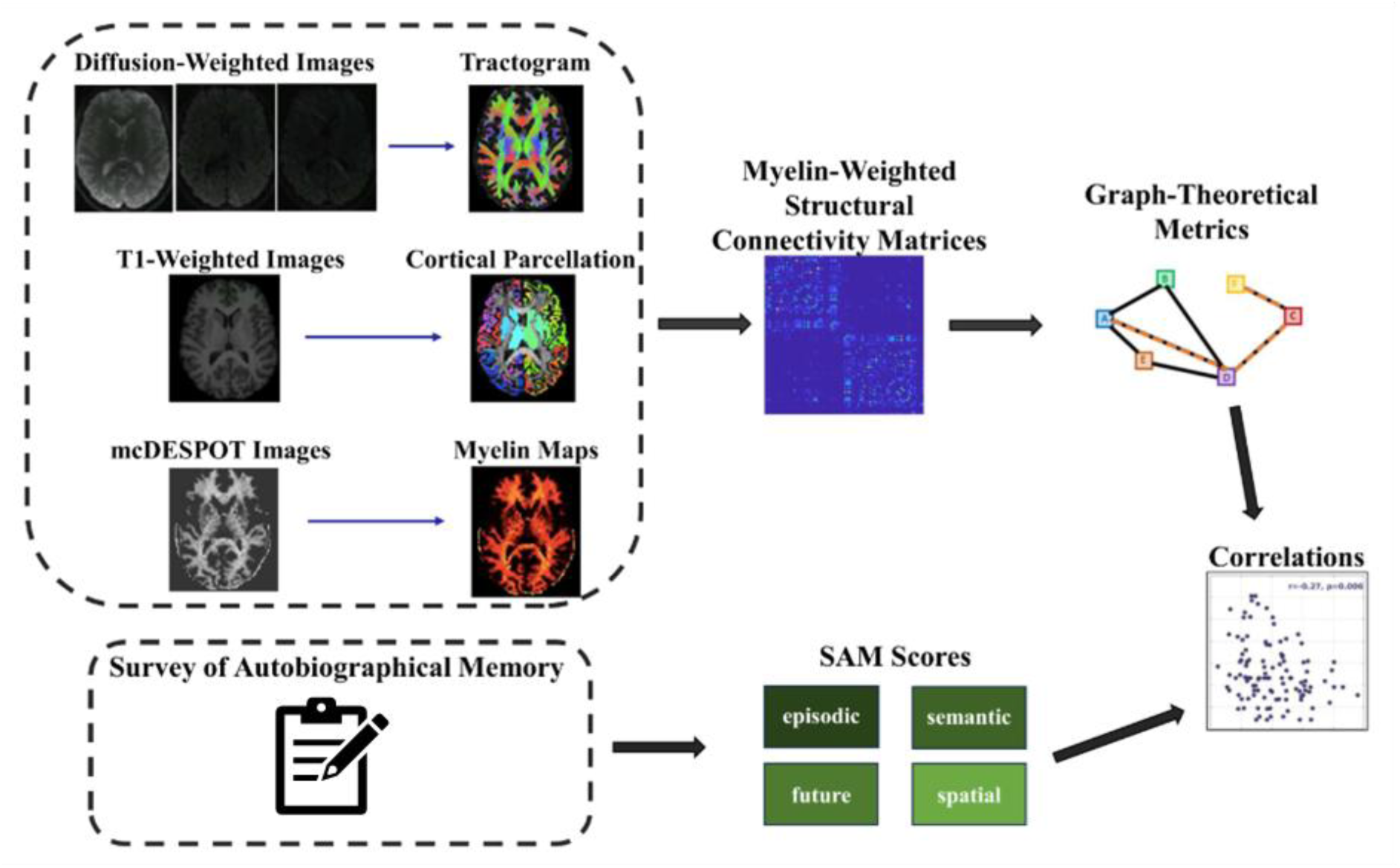
Analysis pipeline.

### 2.1 Participants

The WAND study recruited 170 healthy participants between 18 and 63 years of age. Exclusion criteria for taking part were: diagnosis of any heart or breathing problem; high blood pressure; nerve issues, including carpal tunnel syndrome and nerve damage; history of stroke, brain tumour or brain injury; dizziness, palpitations or fainting; diabetes; current or previous diagnosis of psychiatric condition; use of medication known to alter breathing, blood pressure or mood; pregnancy or breast-feeding; heavy use of tobacco; frequent migraines; epilepsy; and history of concussion resulting in loss of consciousness. The study was approved by the Cardiff University School of Psychology Research Ethics Committee (EC.18.08.14.5332RA3). All participants gave informed written consent and permission for their anonymised data to be shared with researchers in other organisations via a publicly accessible database.

### 2.2 Survey of Autobiographical Memory (SAM)

The SAM was used to measure self-estimated autobiographical memory abilities (Palombo et al., 2013) and was administered as part of a wider battery of cognitive tasks and self-report instruments (McNabb et al., 2025). The SAM asks participants to rate their general mnemonic abilities according to various statements. It is divided into four subcategories, referring to a participant’s self-reported capacity for episodic (8 items), semantic (6 items), and spatial memory (6 items), as well as their ability to imagine possible future events (6 items). These are rated on a five-point Likert scale from ‘strongly disagree’ to ‘strongly agree’. SAM-episodic, SAM-semantic, SAM-spatial and SAM-future scores were calculated according to the original weighting protocol courtesy of Brian Levine, in which responses are weighted and calculated together to provide an average score that centres around 100 (*n.b.* these scores differ from those published in McNabb et al., 2025 which were instead computed by averaging each subscale).

### 2.3 MRI data collection

MRI data were collected on a Siemens Connectom 3T scanner with ultra-strong (300mT/m) magnetic field gradients (Siemens Healthcare, Erlangen, Germany). Details can be found in the data release publication (McNabb et al., 2025) but we outline the data collection protocols here for completeness.

T1-weighted images were acquired using magnetization-prepared 180° radio-frequency pulses and rapid gradient-echo (MPRAGE) sequence with repetition time (TR) 2300ms, echo time (TE) 2ms, field of view (FOV) 256 x 256 x 192 mm, matrix size 256 x 256 x 192, voxel size 1 x 1 x 1 mm, flip angle 9°, inversion time (TI) 857ms, in-plane acceleration (GRAPPA) factor 2 and phase-encoding direction anterior to posterior (A>>P).

Multi-shell diffusion-weighted MRI data were acquired using a single-shot spin-echo, echo-planar imaging sequence, in anterior to posterior (A>>P) and posterior to anterior (P>>A) phase-encoding directions. A>>P data comprised two shells of 20 diffusion encoding directions uniformly distributed (Jones et al., 1999) at b = 200 s/mm^2^ and b = 500 s/mm^2^, one shell of 30 directions at b = 1200 s/mm^2^ and three shells of 61 directions each at b = 2400 s/mm^2^, 4000 s/mm^2^ and 6000 s/mm^2^. Two leading non-diffusion-weighted (b = 0 s/mm^2^) images and 11 non-diffusion-weighted images were acquired, dispersed throughout (33^rd^ volume and every 20^th^ volume thereafter). P>>A data comprised two leading non-diffusion-weighted images, one shell of 30 directions at b = 1200 s/mm^2^ and a final non-diffusion-weighted image. Data acquisition details for all b-values are as follows: TR 3000ms, TE 59ms, FOV 220 x 200 mm in-plane, matrix size 110 x 110 x 66, voxel size 2 x 2 x 2 mm, with in-plane acceleration (GRAPPA) factor 2. The diffusion gradient duration and separation were 7ms and 24ms, respectively.

Multicomponent driven equilibrium single pulse observation of T1 and T2 (mcDESPOT) data were acquired (Deoni et al., 2013), using a T1-weighted 3D spoiled gradient-recalled echo sequence (SPGR), an inversion recovery-prepped spoiled gradient-recalled echo sequence (SPGR-IR) and a steady-state free precession (SSFP) sequence. For SPGR: TR=4ms, TE=1.9ms, 8 flip angles (3, 4, 5, 6, 7, 9, 13 and 18*°*); for SPGR-IR: TR=4ms, TE=1.9ms, flip angle 5*°*, full k-space acquisition in PE and slice directions; for SSFP: TR=4.54ms, TE=2.27ms, 8 flip angles (10.00, 13.33, 16.67, 20.00, 23.33, 30.00, 43.33 and 60.00*°*). The phase-encoding direction was A>>P, FOV 220 × 220 × 179 mm^3^, matrix size 128 × 128 × 104 and voxel size 1.72 × 1.72 × 1.72 mm^3^.

The preprocessing and analysis of the diffusion data was performed as described in recent publications (Cai et al., 2025; Messaritaki et al., 2023, 2025) and are included here for completeness.

### 2.4 Parcellation of cortical and subcortical areas

Cortical reconstruction and volumetric segmentation of the T1-weighted images was performed using recon-all from the Freesurfer image analysis suite (https://surfer.nmr.mgh.harvard.edu) (Dale & Sereno, 1993; Dale et al., 1999; B Fischl & Dale, 2000; Fischl et al., 2002; B Fischl et al., 2001, 2004; B Fischl, Sereno, & Dale, 1999; B Fischl, Sereno, Tootell, et al., 1999; Han et al., 2006; Jovicich et al., 2006; Reuter et al., 2010, 2012; Ségonne et al., 2004). The Desikan-Killiany (DK) atlas (Desikan et al., 2006) was used to identify the grey matter areas that correspond to the nodes of the ‘core network’ (a subset of default network regions) (Benoit & Schacter, 2015; Beaty & Schacter, 2018). The DK atlas identifies 82 cortical and subcortical areas and two areas for the left and right cerebellum. The parcellation reconstructed for each participant was overlayed on the T1-weighted image and was visually inspected to ensure that there was good alignment between the two.

### 2.5 MRI Data Pre-processing

Data were analyzed on a CentOS Linux 7 cluster computing system. The diffusion-weighted MRI data were corrected for thermal noise, signal drift, susceptibility-induced distortions, motion and eddy current-induced distortions, gradient non-linearity, and Gibbs ringing artifacts using in-house pipelines and publicly available software. Brain extraction was performed using FSL 6.0.3 (Jenkinson et al., 2012; S M Smith, 2002) and all non-brain data were excluded. Diffusion MRI noise level estimation and denoising was performed using a Marchenko-Pastur principal component analysis (MP-PCA)-based approach (Veraart, Fieremans, et al., 2016; Veraart, Novikov, et al., 2016). Within-image intensity drift was corrected by fitting the diffusion-weighted MRI data to temporally interspersed non-diffusion-weighted images using inhouse code in MATLAB R2017b (MathWorks Inc. Natick, Massachusetts, USA). Slicewise OutLIer Detection (SOLID, (Sairanen et al., 2018)) was applied, employing lower and upper thresholds of 3.5 and 10 respectively, using a modified Z-score and a variance-based intensity metric. The susceptibility-induced off-resonance field was estimated from the non-diffusion-weighted data collected in opposing (A>>P and P>>A) phase-encoding directions using FSL’s topup function (Andersson et al., 2003; Stephen M Smith et al., 2004) and corrected, along with eddy current-induced distortions and subject movements, using FSL’s eddy tool (Andersson & Sotiropoulos, 2016). In-house code was used to correct for gradient non-uniformity distortions in MATLAB R2017b (MathWorks Inc. Natick, Massachusetts, USA). Gibbs ringing correction was performed in MRtrix3 using the subvoxel-shifts method (Kellner et al., 2016). The preprocessed images were checked to ascertain lack of structure in noise maps, absence of significant slice-wise outliers, reduction of movement, good alignment between volumes, lack of implausible signal values and successful gradient non-uniformity correction. Any data that did not satisfy those criteria were excluded from analysis. The diffusion tensor (Pierpaoli et al., 1996) was calculated from the diffusion-weighted images (DWIs).

Preprocessing of mcDESPOT data included brain extraction of T1-weighted MPRAGE data using HD-BET (Isensee et al., 2019) followed by resampling of MPRAGE and masked brain-extracted data to SPGR-IR resolution using Advanced Normalization Tools (Avants et al., 2009). SSFP and SPGR data were corrected for motion using the FMRIB Software Library (FSL version 6.0.1) motion correction tool (Jenkinson et al., 2002) and then SPGR-IR data were registered to the motion corrected SPGR data using FSL’s linear registration tool (Jenkinson & Smith, 2001). These data were then used to create B1, T1 and B0 maps using functions from QUantitative Imaging Tools (QUIT; (C Wood, 2018)), including driven equilibrium single pulse observation of T1 (DESPOT1) with and without high-speed incorporation, as well as DESPOT2 (Deoni, 2007; Deoni et al., 2003, 2008). Finally, a three-compartment model (Deoni et al., 2013) was used to separate SPGR and SSFP signals into discrete pools representing myelin water and intra/extra-cellular water or cerebrospinal fluid.

### 2.6 Calculation of fibre orientation distributions and response functions, and tractography

MRtrix3 version 3.0.3 (Tournier et al., 2019) was used to calculate the response function and the fibre orientations, and to perform anatomically-constrained tractography. The response function was calculated using the Dhollander algorithm (Dhollander et al., 2016, 2019). Fibre orientation distributions were calculated using the Multi-Shell-Multi-Tissue constrained spherical deconvolution algorithm (Jeurissen et al., 2014) which allows estimation of multiple crossing fibres within each voxel.

MRtrix3 was used to segment anatomical images (R. E. Smith et al., 2012; Stephen M Smith et al., 2004; S M Smith, 2002) into cortical/subcortical grey matter, cerebrospinal fluid, white matter, and abnormalities. Anatomical images were co-registered to the DWIs using the flirt algorithm of FSL with the nearest neighbour option (Jenkinson et al., 2012). The interface between grey and white matter was identified using MRTrix3 (Tournier et al., 2019). Anatomically-constrained streamline tractography (R. E. Smith et al., 2012) was used to generate whole-brain tractograms with the seeds in the grey-white matter interface. The minimum and maximum streamline lengths were 30 and 250mm respectively, the cutoff was 0.06, and the maximum angle between successive steps was 50 degrees. Twenty million streamlines were generated for each participant. The SIFT2 algorithm (R. E. Smith et al., 2015) was used to provide tractograms that have a density of reconstructed connections proportional to the fibre density within each voxel as estimated by the diffusion model. The tractogram reconstructed for each participant was overlayed onto the participant’s fractional anisotropy map and visually inspected to make sure that there was good coverage of the white matter areas and no streamlines extended into unphysical areas.

### 2.7 Network generation and edge weight choice

The structural brain networks were constructed by combining the results from the cortical parcellation with those from the tractography. The nodes were the cortical and subcortical brain areas as identified through the Desikan-Killiany atlas, and the edges (connections) were the white matter tracts connecting those areas.

The mcDESPOT data allowed us to calculate the myelin water fraction (MWF), i.e., the fraction of the T2 signal that comes from the water trapped between the lipid bilayers of the myelin sheath, in the voxels along the white matter tracts. We use the mean MWF as edge weight for the structural brain networks, taking the mean MWF across the streamlines for each tract.

We investigated correlations of the SAM subscale scores with the topological measures of the core memory network. To define the network, we chose the Desikan-Killiany brain areas shown to co-activate and covary consistently during mnemonic cognition and largely overlap with the DMN (Andrews-Hanna, 2012; Benoit & Schacter, 2015; Beaty & Schacter, 2018; Braga, 2025; Girn et al., 2024; Menon, 2023; Li et al., 2021; Sylvester et al., 2020). The ROIs sampled: parahippocampal gyrus, isthmus cingulate, precuneus, inferior parietal lobe, posterior cingulate cortex, medial orbitofrontal cortex, rostral anterior cingulate, lateral orbitofrontal, caudal anterior cingulate, thalamus, hippocampus, amygdala, temporal pole. See Fig. 2.

**Figure 2:**
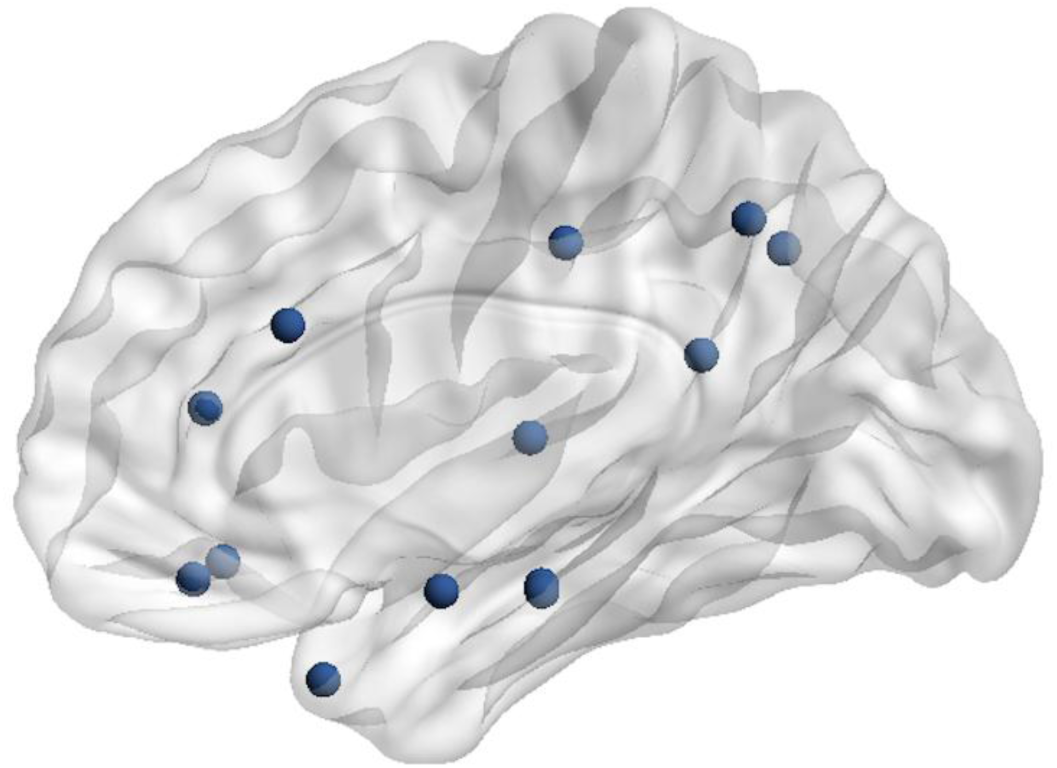
Visualisation of core network regions of interest. Image prepared using BrainNet Viewer (https://www.nitrc.org/projects/bnv)

### 2.8 Graph theoretical metrics

We used the Brain Connectivity Toolbox (Rubinov & Sporns, 2010) to calculate the mean nodal strength, mean clustering coefficient, characteristic path length and global efficiency for the core network. These measures constitute a set of weighted graph theoretical metrics that provide information on network integration and segregation (Petersen & Sporns, 2015). To avoid multi-collinearity in our analysis, we calculated the Pearson correlation coefficient of each of these metrics with the others. If two graph theoretical metrics were highly correlated with each other (i.e., correlation coefficient of 0.9 or higher) we kept only one of the two.

We used the Mahalanobis distance to identify multivariate outliers, and excluded them from further analysis (Hadi et al., 2009). We then calculated the Spearman partial correlation coefficients, controlling for participant age and sex, between each of the selected graph theoretical metrics of the core network and each of the 4 SAM subscale scores. We used the false-discovery-rate algorithm to correct for multiple comparisons.

To test whether the graph theoretical metrics of the core network are preferentially associated with SAM subscale scores (as opposed to other subnetworks with the same number of nodes), we randomly selected 10,000 subnetworks with the same number of brain areas as the core network. We calculated the same graph theoretical metrics and repeated the age- and sex-corrected Spearman correlation analysis for each of those randomly generated subnetworks. This resulted in a distribution of values for each Spearman partial correlation coefficient. The correlation coefficients calculated for the core network analysis were plotted against those distributions to evaluate the likelihood that the observed coefficients could have resulted by chance if any random subnetwork were being used.

To understand whether any statistically significant correlations for the non-global graph theoretical metrics (i.e., mean clustering coefficient and mean nodal strength) are driven by specific nodes of the core network or by all of its nodes, we performed a node-specific analysis. We calculated the age- and sex-corrected Spearman partial correlation coefficients between the graph theoretical metric in question for each node of the core network and the corresponding SAM score.

All analyses were run in in MATLAB R2021b (MathWorks Inc. Natick, Massachusetts, USA).

## 3. Results

### 3.1 Participants

There were 109 participants (67 female, age mean 29.7 years, age SD 10.4 years) with diffusion MRI and mcDESPOT data who completed the SAM (distribution shown in Fig. 3). A sensitivity analysis conducted using G*Power (Version 3.1.9.7; Faul et al., 2007) indicated that the final sample of n= 109 had 80% power to detect a two-tailed medium-sized (Gignac & Szodorai, 2016) correlation of 0.26 at p < .05.

**Figure 3:**
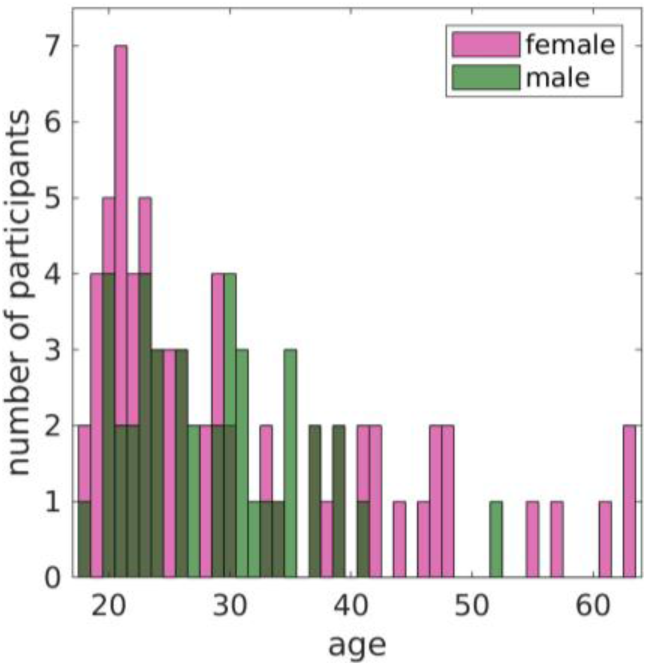
Age distribution of the participants.

### 3.2 SAM data quality

The mean, standard deviation and range for all subscales (SAM-episodic: mean= 97.8, SD= 13.8, range= 73-137; SAM-semantic: mean= 97.2, SD= 12.6, range = 75.6-135; SAM-spatial: mean= 98.7, SD= 14.3, range= 68.6-121; SAM-future: mean= 91.2, SD= 8.57 range= 78-113) were comparable to that of previous studies (Setton et al., 2022) and the internal consistencies of all the subscales were α > .63 (SAM-episodic= .83; SAM-future= .63; SAM-semantic= .65; SAM-spatial= .76), demonstrating sufficient inter-item reliability (Nunnally, 1978). SAM-episodic was significantly correlated with SAM-semantic (ρ(107)= .56, p< .001; Fig. 4) and SAM-spatial (ρ(107)= .32, p< .001; Fig. 4). Significant correlations were found between the SAM-semantic and SAM-future subscales (ρ(107)= .23, p= .017; Fig. 4). The episodic and semantic scores were most strongly correlated, replicating previous findings (Palombo et al., 2013), showing that the SAM demonstrates reliability as a self-report measure of perceived mnemonic capacity. Sex differences were also examined: males (mean= 103, SD= 13) scored significantly higher than females (mean= 96.1, SD = 14.5) on SAM-spatial only (t(104)= -2.62, p= .01, d = -0.53), which aligns with previous findings (Palombo et al., 2013; Setton et al., 2022). No significant correlations were found with age for SAM-episodic (r= .07, p= .42), SAM-semantic (r= -.01, p= .88), SAM-spatial (r= .15, p= .11) or SAM-future (r= -.06, p= .55). Jamovi (Version 2.5.6; The jamovi project, 2023) was used to calculate these means, standard deviations, ranges, Cronbach’s alphas, correlations, and t-tests.

**Figure 4:**
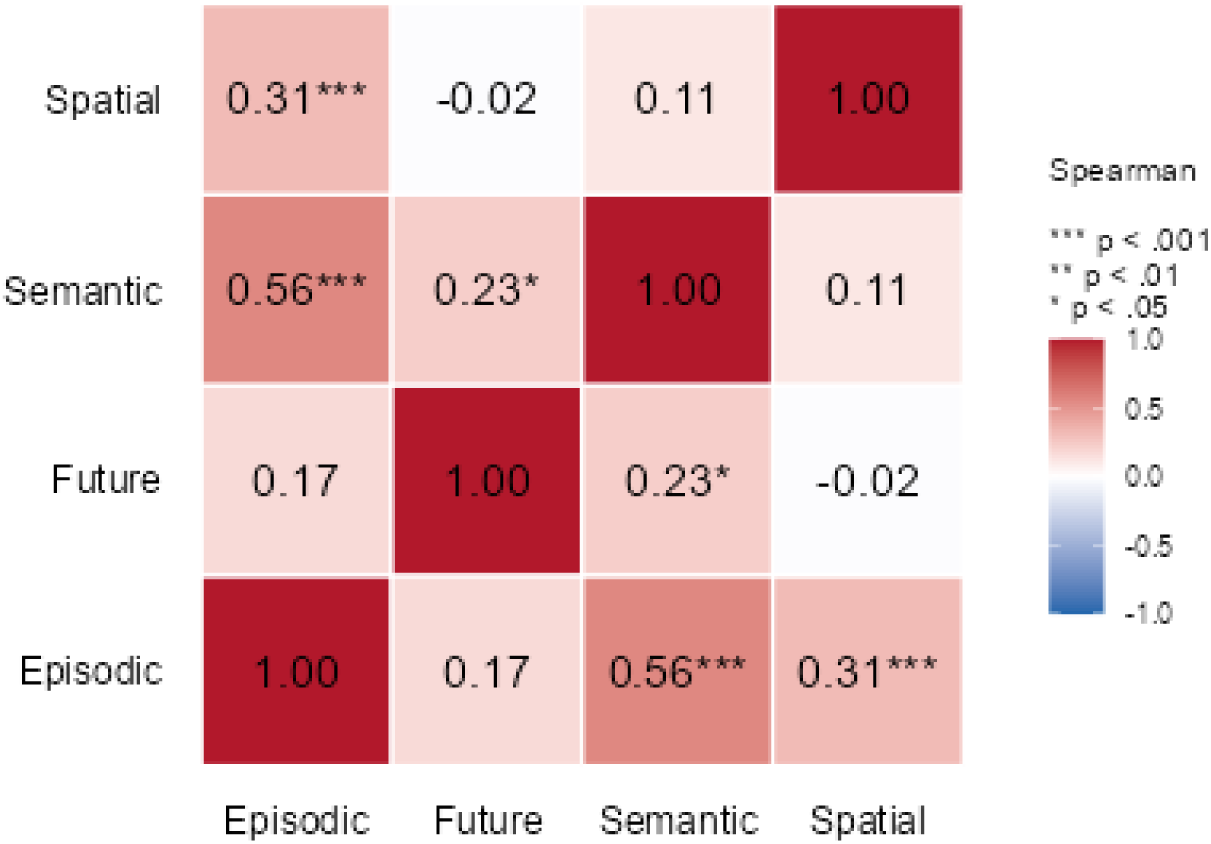
Spearman correlations between SAM subscale scores.

### 3.3 Correlation analysis for the graph theoretical metrics

The results of the Pearson correlation analysis for the graph theoretical metrics are shown in Table 1. The global efficiency of the core network correlated strongly with the characteristic path length, and the characteristic path length correlated strongly with the mean nodal strength. We kept the characteristic path length and the mean clustering coefficient of the networks as the graph theoretical metrics for our analysis.

**Table 1:** Correlation coefficients between the graph theory metrics.

|  | Global Efficiency | Mean Clustering Coefficient | Mean Nodal Strength |
| --- | --- | --- | --- |
| Characteristic Path Length | -0.98 | -0.85 | -0.97 |
| **Global Efficiency** |  | **0.81** | **0.92** |
| **Mean Clustering Coefficient** |  |  | **0.87** |

### 3.4 Associations between SAM scores and graph theoretical metrics

We observed statistically significant relationships between the SAM-future scores and both the characteristic path length (negative correlation) and the mean clustering coefficient (positive correlation) of the core network. The former relationship survived FDR multiple comparison correction, while the latter did not. The effect sizes were medium for both the significant correlations (Gignac & Szodorai, 2016). All age- and sex-corrected Spearman correlation coefficients and *p*-values are shown in Table 2. The scatter plots for the nominally significant correlations (i.e., p-value < 0.05) are shown in Fig. 5.

**Table 2:** Spearman correlation coefficients and *p*-values between the SAM scores and the graph theoretical metrics of the core network (age- and sex-corrected). Bold font indicates statistically significant correlations after FDR multiple comparison correction, while the underlined values are nominally significant (p<0.05) but do not survive FDR correction.

|  | **SAM-Episodic** | **SAM- Semantic** | **SAM-Spatial** | **SAM-Future** |
| --- | --- | --- | --- | --- |
| **Characteristic Path Length** | $r = 0.08$<br>$p = 0.39$ | $r = -0.02$<br>$p = 0.84$ | $r = -0.02$<br>$p = 0.85$ | **$r = -0.27$**<br>**$p = 0.006$** |
| **Mean Clustering Coefficient** | $r = -0.11$<br>$p = 0.23$ | $r = 0.08$<br>$p = 0.41$ | $r = 0.06$<br>$p = 0.52$ | <u>$r = 0.23$</u><br><u>$p = 0.019$</u> |

**Figure 5:**
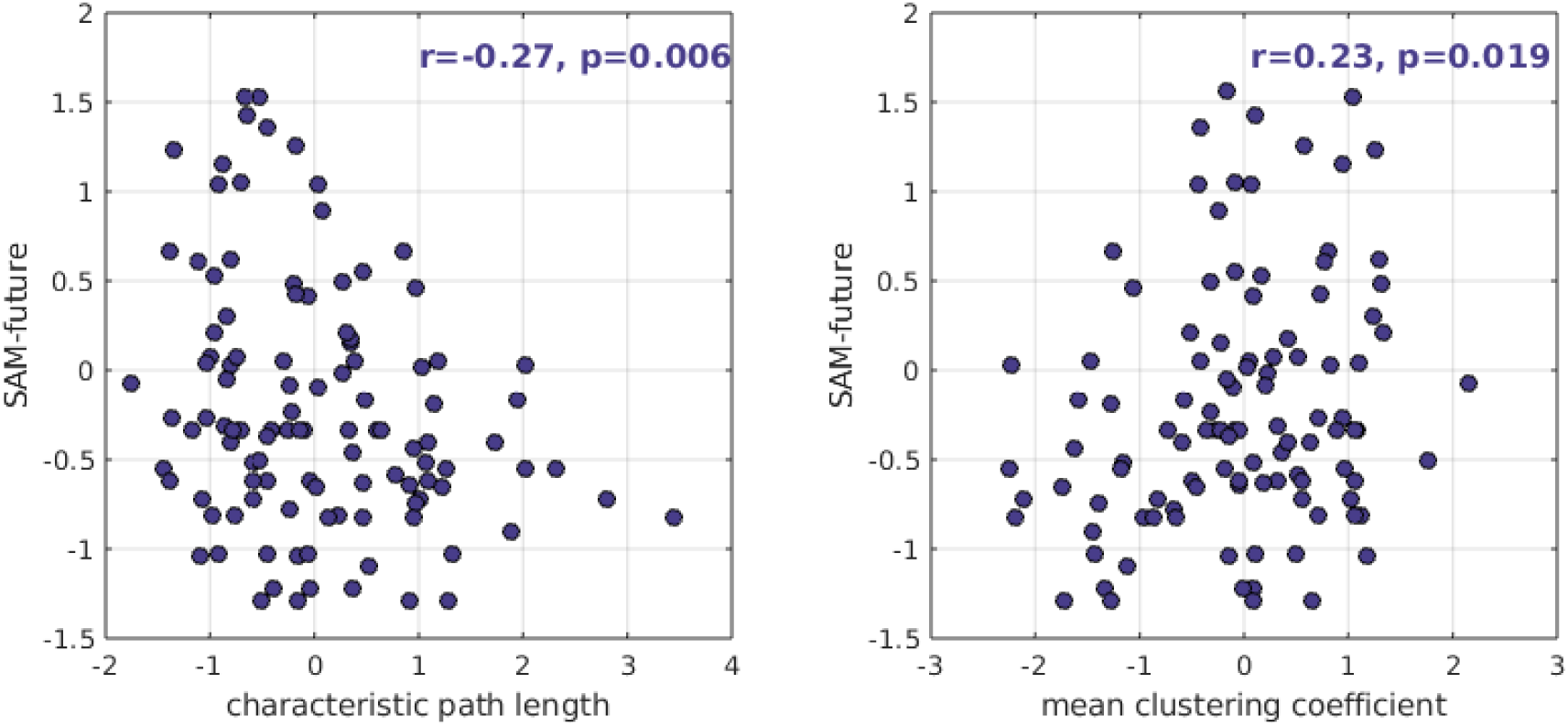
Scatter plots for the SAM scores vs the graph theoretical metrics of the MWF-weighted core network, for the age- and sex-corrected Spearman correlations that are nominally statistically significant.

### 3.5 Control analysis

The analysis of the 10,000 non-task-specific networks showed that the results are highly likely to be specific to the core memory network rather than the reduction of nodes in it in comparison to the whole brain network (Fig. 6). The observed correlation between SAM-future score and characteristic path length was stronger than 97.8% of the values for the non-task-specific subnetworks (2.01 SD away from the mean, *p*=0.022). The observed correlation between SAM-future and mean clustering coefficient was stronger than 86.1% of the values for the non-task-specific subnetworks (1.10 SD away from the mean, *p* = 0.135).

**Figure 6:**
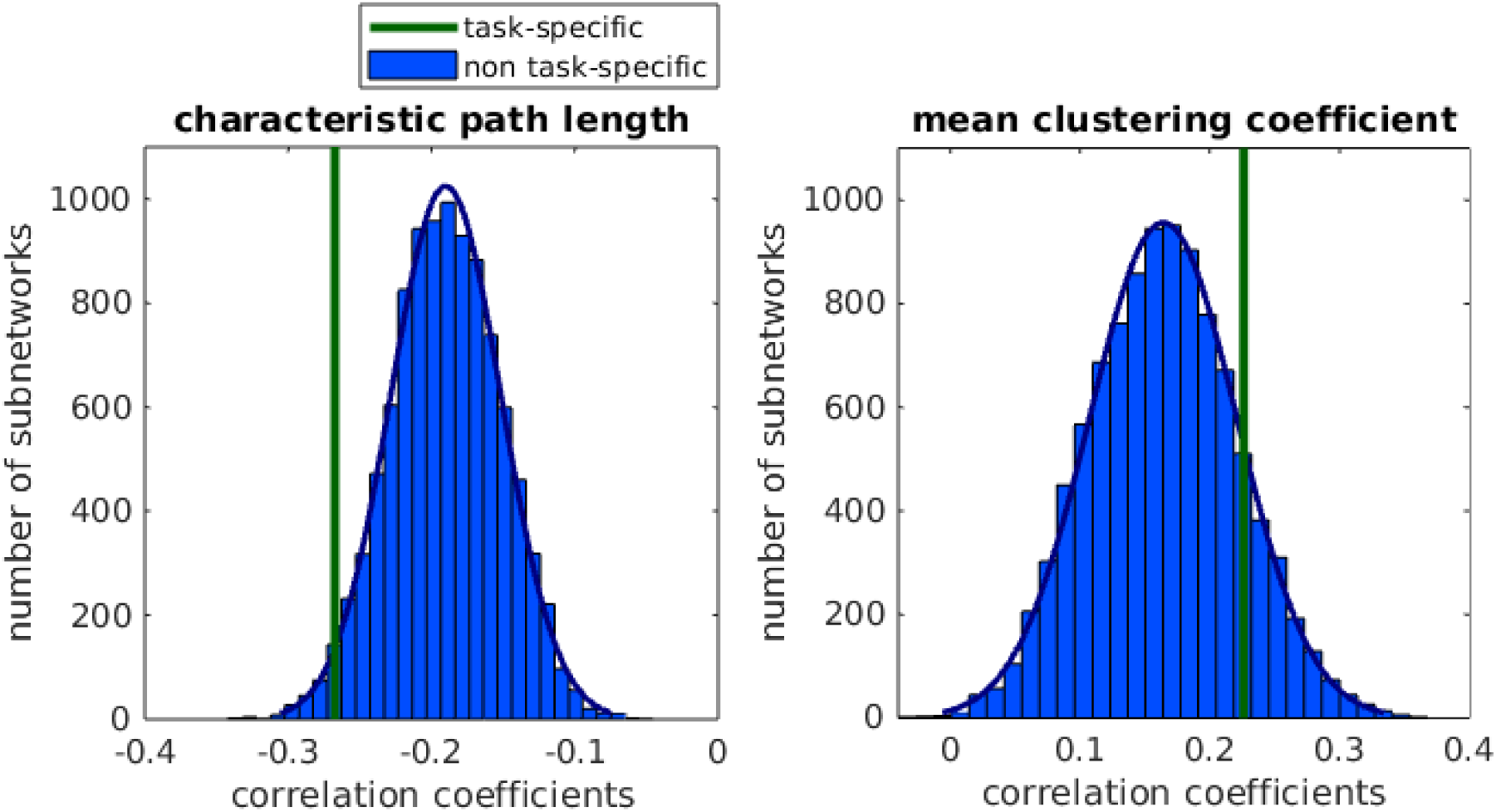
Distributions of age- and sex-corrected Spearman correlation coefficients of the non-task-specific subnetwork, for the correlation coefficients that are nominally statistically significant in the analysis for the task-specific subnetwork. The correlation coefficients measured for the core network are indicated by the vertical lines.

### 3.6 Node-specific analysis

Table 3 and Fig. 7A show the age- and sex-corrected Spearman correlation coefficients between the SAM-future score and the clustering coefficient for each node of the core network. All nodes exhibited positive correlations, in agreement with the positive relationship between mean clustering coefficient and SAM-future score for the network. The ROIs that survived FDR multiple comparison correction were the bilateral caudal anterior cingulate, lateral orbitofrontal, medial orbitofrontal, rostral anterior cingulate, temporal pole, and amygdala as well as the left isthmus cingulate, precuneus and hippocampus. The effect sizes of these relationships ranged from medium to large (Gignac & Szodorai, 2016). A Wilcoxon signed-rank test comparing correlation coefficients between the left and right hemispheres showed correlations were significantly higher in the left hemisphere than the right hemisphere (*z*= 2.94; *p*= .004; Fig. 7B), consistent with evidence of a modest left hemisphere bias in autobiographical memory (Daviddi et al., 2023; Daviddi et al., 2025).

**Table 3:** Node-specific correlation coefficients and *p*-values. The bold-typed values are the ones that survive FDR multiple comparison correction.

| ROI | SAM-future score vs node clustering coefficient |  |  |  |
| --- | --- | --- | --- | --- |
|  | Left hemisphere |  | Right hemisphere |  |
| | Correlation ($p$-value) | Nodal degree averaged over participants | Correlation ($p$-value) | Nodal degree averaged over participants |
| Caudal anterior cingulate | **0.22 (0.023)** | 22.8 | **0.22 (0.024)** | 22.9 |
| Inferior parietal | 0.20 (0.045) | 24.5 | 0.17 (0.086) | 24.7 |
| Isthmus cingulate | **0.23 (0.018)** | 24.7 | 0.18 (0.065) | 24.8 |
| Lateral orbitofrontal | **0.31 (0.001)** | 23.3 | **0.25 (0.010)** | 23.5 |
| Medial orbitofrontal | **0.26 (0.007)** | 23.1 | **0.27 (0.005)** | 22.8 |
| Parahippocampal | 0.19 (0.054) | 20.9 | 0.16 (0.107) | 21.3 |
| Posterior cingulate | 0.20 (0.045) | 23.9 | 0.19 (0.051) | 23.8 |
| Precuneus | **0.23 (0.014)** | 24.9 | 0.19 (0.057) | 24.9 |
| Rostral anterior cingulate | **0.30 (0.002)** | 22.3 | **0.26 (0.006)** | 22.4 |
| Temporal pole | **0.31 (0.001)** | 20.1 | **0.22 (0.026)** | 17.4 |
| Thalamus | 0.20 (0.043) | 24.7 | 0.17 (0.073) | 24.9 |
| Hippocampus | **0.23 (0.016)** | 23.9 | 0.21 (0.032) | 24.3 |
| Amygdala | **0.28 (0.004)** | 20.8 | **0.26 (0.006)** | 21.1 |

**Figure 7.**
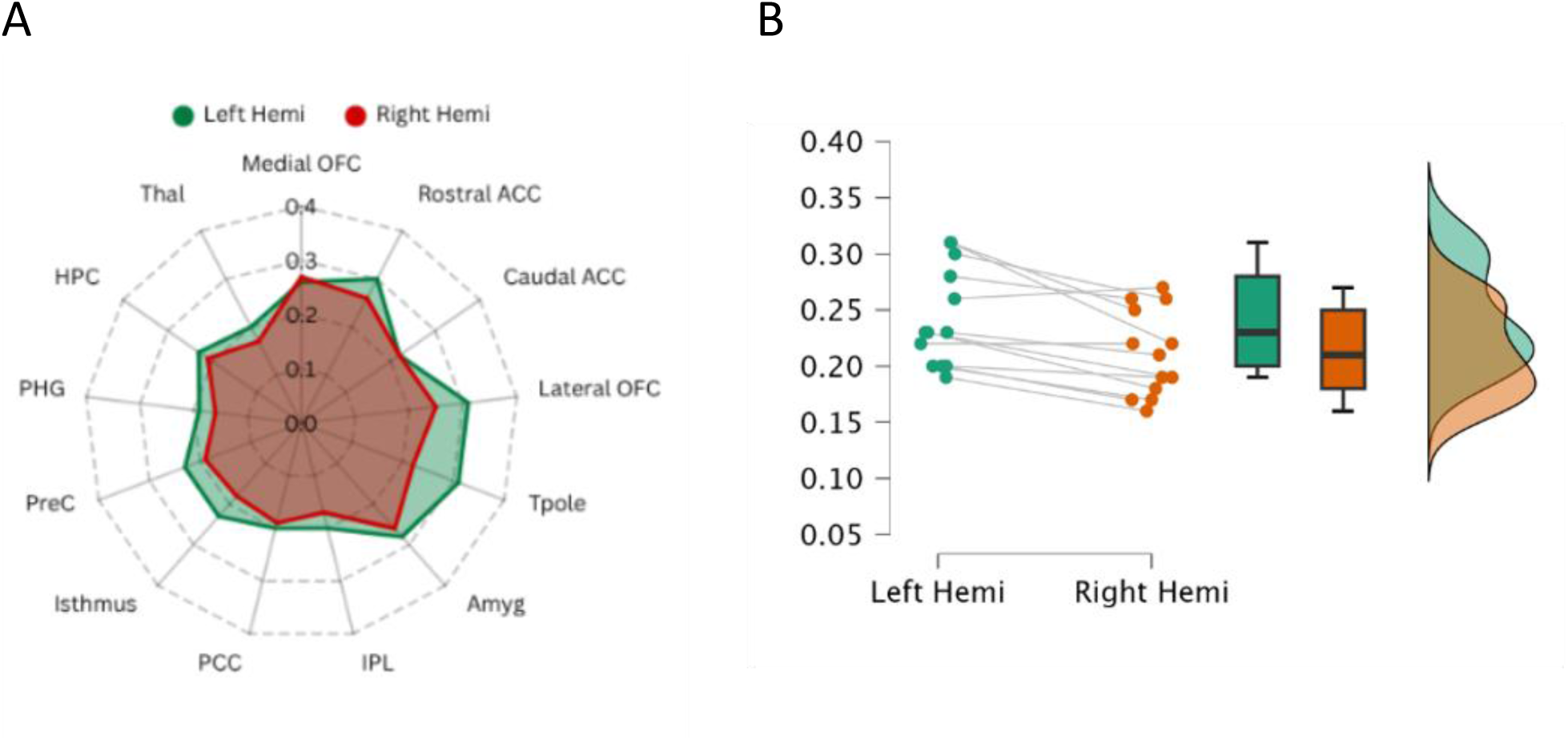
A) Radar plot depicting the node-specific correlation coefficients B) Raincloud plot depicting correlation coefficients for left and right hemisphere regions.

Further analysis showed that there is a strong relationship between the average (over participants) of the node degree of a node and the strength of correlation of its clustering coefficient with the SAM-future score (Fig. 8; Spearman correlation: -0.46, *p*=0.018). We elaborate on this in the Discussion.

**Figure 8:**
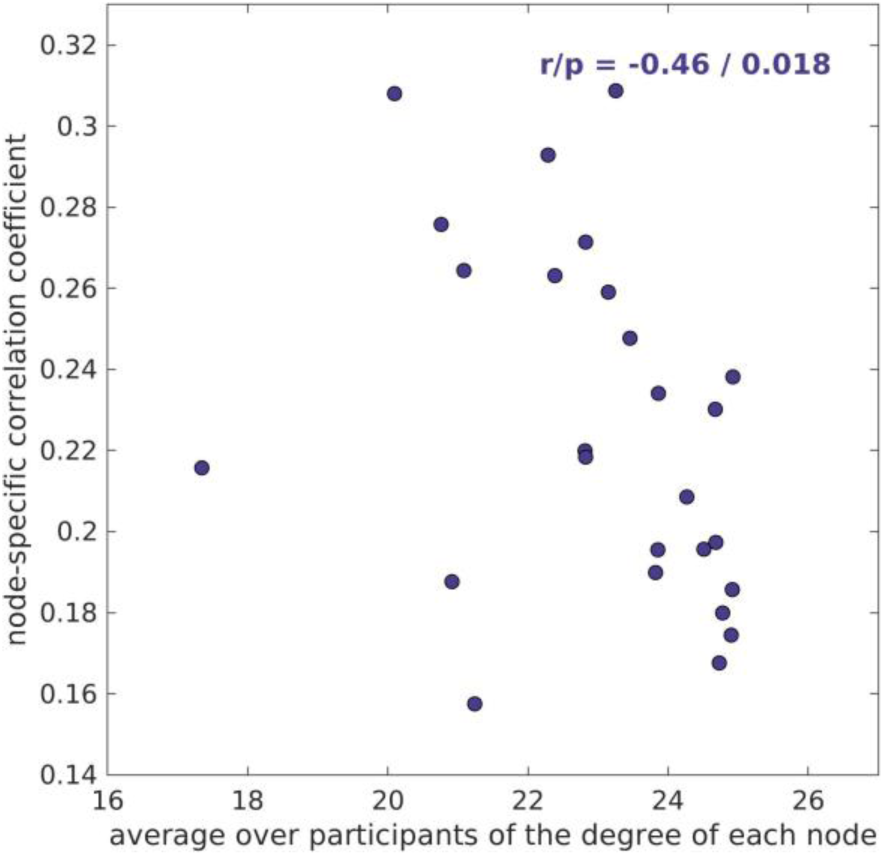
Relationship between the average (over participants) nodal degree and the node-specific correlation coefficients between SAM-future score and mean clustering coefficient. The Spearman correlation coefficient and p-value are shown.

## 4. Discussion

We investigated the relationship between individual differences in trait autobiographical memory and the organisational properties of the core network in a large cohort of healthy participants. This is the first study to examine the role of myelin-weighted structural brain networks in mnemonic cognition. Myelin is critical for regulating neural signalling and synchronisation across distributed circuits (Bells et al., 2017; Drakesmith et al., 2019; Suminaite et al., 2019). MRI-based biomarkers of myelin, including MWF, have been associated with electrophysiological functional connectivity (Cai et al., 2025; Messaritaki et al., 2021), autobiographical memory (Clark et al., 2022), and large-scale structure-function coupling (Nelson et al., 2026). Unlike fractional anisotropy of the diffusion tensor and streamline count, which are influenced by factors including axonal density and directionality (Jones et al., 2013), MWF is more directly related to myelin (Bae et al., 2026).

### 4.1 SAM-future and graph theoretical metrics

We found that SAM-future scores were negatively correlated with characteristic path length and positively correlated with mean clustering coefficient of the core network. No significant relationships were found for the episodic, semantic or spatial subscales.

No prior research has elucidated the neural correlates of SAM-future scores: previous functional imaging studies using the SAM found no significant associations with this subscale (Petrican et al., 2020; Teghil et al., 2022). Our findings support our hypothesis and align with resting-state fMRI studies demonstrating the involvement of the DMN in future thinking to a greater extent than episodic recollection (Andrews-Hanna et al., 2010; Bellana et al., 2017; Gilmore et al., 2018; Kucyi & Davis, 2014; Preminger et al., 2011; Ostby et al., 2012; Villena-Gonzalez et al., 2018). The observed pattern suggests a core network that combines global integration (reflected by shorter path lengths) with local specialisation (reflected by higher clustering), consistent with an efficient small world topology (Bullmore & Sporns, 2012; Rubinov & Sporns, 2010).

One question is why our analysis showed correlations between graph theoretical metrics and SAM-future only, when all SAM domains are linked to the core network (Menon, 2023). Firstly, the SAM-future subscale may have higher construct validity than the other subscales, as indexed by its factor structure (Picco et al., 2020; Setton et al., 2022). Secondly, sensitivity analyses indicated statistical power to detect medium effect sizes and therefore, our sample size may have been insufficient to detect lower effect sizes possibly present in the other subscales. However, our findings align with the constructive episodic simulation and scene construction hypotheses (Hassabis & Maguire, 2007; Schacter & Addis, 2007) which posit that future thinking places greater demands on the core network in comparison to memory, as the former depends heavily on the flexible recombination of episodic information into novel representations, whereas the latter requires reactivation of an existing representation. Our results add nuance to this theory, suggesting that integration and segregation within the core network may be particularly relevant for constructing future events rather than memory-related processes (e.g. encoding, retrieval). Other networks (e.g. visual; Petrican et al., 2020) or DMN subnetworks (e.g. MTL; Sheldon et al., 2016) may be more relevant to individual differences in memory. Indeed, future thinking may rely more on core network function than episodic recall (Benoit & Schacter, 2015; Xu et al., 2016) and link more to imaginative traits (e.g. openness; Beaty et al., 2016) associated with wider DMN function. This aligns with the original SAM validation study, which found only a modest association between SAM-episodic and SAM-future (Palombo et al., 2013). Based on this result, the authors proposed self-reported future thinking depends not only on episodic memory, but other cognitive abilities such as problem solving and social reasoning. Taken with evidence that autobiographical memory and future thinking exhibit differences and dissociations (Schacter et al., 2012), this indicates that future thinking depends on additional mechanisms beyond episodic memory.

### 4.2 Node-specific analysis

The post-hoc analysis of the clustering coefficient identified the core network nodes driving the association between local clustering and SAM-future scores. Individuals with more integrated subnetworks in these nodes may better coordinate the multiple cognitive processes required to simulate future events. Because several of these nodes also support episodic and semantic memory, we highlight that the absence of associations with other SAM subscales does not preclude possible node-specific relationships.

It is notable that relationships with the left hippocampus, isthmus cingulate and bilateral temporal poles were significant, because these regions are linked with the reconstructive processes of future thinking. The hippocampus supports the flexible recombination of episodic details (Schacter & Addis, 2007, 2008) and the construction of spatially coherent scenes (Hassabis et al., 2007; Hassabis & Maguire, 2007). Indeed, future thinking is reliably associated with heightened hippocampal activation (Schacter et al., 2012), and the magnitude of this activity scales with the amount of episodic detail generated (Hassabis et al., 2007; McLelland et al., 2015). Similarly, the involvement of the isthmus cingulate, which links the retrosplenial/posterior cingulate gyrus to the parahippocampal gyrus, is consistent with work demonstrating that the parahippacampal and retrosplenial cortices are implicated in contextual imagery processing (Bar & Aminoff, 2003), scene construction (Vann et al., 2009), and detail elaboration (Addis et al., 2007). Greater clustering within the isthmus cingulate may therefore facilitate efficient information exchange between these regions, supporting the generation of richer future events. On the other hand, the temporal pole is a key hub for schematic knowledge (Visser et al., 2010) which plays a crucial role in imagining the future (Fivush, 2011; Gilboa & Marlatte, 2017; Irish & Piguet, 2013; Levine et al., 2002; Renoult et al., 2019; Tulving, 2002; Yeshurun et al., 2021). Compared with remembering, future event construction relies more on schemas, because imagining a novel event requires substantial general knowledge (Addis et al., 2025; Gilboa & Marlatte, 2017; Irish & Piguet, 2013; Yeshurun et al., 2021). This is supported by evidence that individuals with semantic dementia show disproportionately impaired future thinking (Irish et al., 2012a; Irish et al., 2012b; Irish & Piguet, 2013; Viard et al., 2014), with deficits correlating with bilateral temporal pole atrophy (Irish et al., 2012b). Similar impairments are also observed in temporal pole epilepsy (Lechowicz et al., 2016; Rayner et al., 2022) and the left temporal pole is preferentially activated in future imagining relative to remembering (Abraham et al., 2008).

One of the strongest correlations was with the bilateral amygdala, a region widely associated with socio-emotional processing (Phelps & LeDoux, 2005; Sergerie et al., 2008; Waymar & Schwabe, 2016). This aligns with evidence linking emotional intensity to autobiographical event vividness (Habermas & Diel, 2013; Kensinger et al., 2011; Kensinger & Fields, 2024; Phelps & Sharot, 2008; Reisberg et al., 1988; Talarico et al., 2004), suggesting that individuals who imagine more emotional future events rate their future thinking as more vivid. This may be specific to SAM-future because it includes items related to vividness and emotionality whereas SAM-episodic focuses on episodic detail and temporal context. Alternatively, this may reflect the amygdala’s broader role in a distributed neural system supporting emotional salience, social cognition and internally generated thought (Crawford et al., 2020; Edmonds et al., 2024; Sylvester et al., 2012; Sylvester et al., 2020). Greater local integration of the amygdala may therefore facilitate emotional and social elaboration of future events.

We observed that nodes with fewer connections showed stronger correlations between clustering coefficient and SAM-future scores, suggesting that nodes with smaller neighbourhoods have more impact on future thinking than those with more extended neighbourhoods. One possibility is that less extensively connected nodes support more specialised aspects of imagining the future, with higher clustering facilitating efficient communication within tightly interconnected subnetworks. While one aspect that could require efficient integration is the flexible recombination of stored episodic content (Schacter & Addis, 2007; Schacter et al., 2017; Schacter & Thakral, 2024), we identified nodes associated with broader cognitive functions including self-referential thought (precuneus; Cavanna & Trimble, 2006), resource allocation (anterior cingulate; Kerns et al., 2004) and value-based evaluation (orbitofrontal cortices; Murray et al., 2015). Local clustering in less extensively connected regions may thus reflect more efficient integration within specialised subnetworks that support social, emotional and episodic processing components. This interpretation is consistent with the modest relationship between SAM-future and SAM-episodic scores (Palombo et al., 2013), suggesting that individual differences in future thinking emerge from coordinated activity across multiple core network subsystems, rather than the episodic memory system alone.

### 4.3. Limitations

The participants in our study were predominantly in the age range of 18 – 30 years old, with only 35% aged 31 – 63 years. As myelin characteristics are known to vary across the lifespan (Ahmad & Yap, 2026; Buyanova & Arsalidou, 2021), our findings may be most relevant to younger adults. As is the case with similar studies, our recruitment methods may be biased in favour of people who are educated for longer than average, thus our results may be missing effects present in the general population.

Setton et al. (2022) has argued that the SAM primarily captures memory confidence and mental imagery-related aspects of remembering. This is supported by evidence that individuals with aphantasia score low on the SAM (Dawes et al., 2020; Sheldon et al., 2016), as well as correlations with object imagery (Fan et al., 2021), scene construction (Clark & Maguire, 2020) and self-efficacy (Setton et al., 2022). This may explain why no relationships were observed with the other subscales, despite evidence linking higher DMN nodal strength to laboratory-based episodic memory performance (Matyi & Spielberg, 2022). This type of performance-based task is less dependent on mental imagery and metacognition than the SAM, and future thinking is more reliant on mental imagery than memory (Conti & Irish, 2021; Dawes et al., 2022). Future research should assess relationships between microstructural metrics and performance-based measures, such as the Autobiographical Interview (Addis et al., 2008; Levine et al., 2002) and scene construction task (Hassabis et al., 2007), to determine whether our results reflect the self-report nature of the SAM, and if these limitations underpin the lack of associations in the other subscales.

## 5. Conclusion

By leveraging high-gradient diffusion MRI to construct myelin-weighted structural brain networks, this work provides a fine-grained account of the structural properties of the core network that support future thinking. Our findings demonstrate that future thinking is supported by a balance of global integration and local clustering within the core network. Future research should establish whether these findings generalise to performance-based measures of future thinking, or whether the observed associations are specific to self-reported, trait-level tendencies. Additionally, examining alternative network parcellations and edge weightings may reveal complementary or distinct correlates, helping to determine the extent to which these findings are uniquely driven by myelin-sensitive measures of brain connectivity. Yet the present findings suggest that individual differences in future thinking may arise not simply from the integrity of isolated brain regions, but from the myelin-sensitive organisation of the broader autobiographical network.

## Acknowledgements

We would like to thank John Evans, Peter Hobden, Ian Driver, Maria Engel, Hannah Thomas, Matthew Ploszajski, Emma Morgan, Svetla Manolova, Vanessa Hyde and Elena Stylianopoulou for contributing to WAND participant recruitment and data collection.

## Open Access

For the purpose of open access, the authors have applied a creative commons attribution (CC BY) licence to any author accepted manuscript version arising. The data and materials for all experiments are available at https://www.nature.com/articles/s41597-024-04154-7#Ack1

## Author Contributions

Lucie S. Reed: Investigation; Conceptualization; Methodology; Formal Analysis; Writing – Original Draft; Writing – Review & Editing; Visualization

Eirini Messaritaki: Investigation; Data Curation; Conceptualization, Methodology; Formal Analysis; Writing – Original Draft; Writing – Review & Editing; Visualization

Craig Hedge: Investigation; Data Curation; Writing – Review & Editing

Carl J. Hodgetts: Investigation; Methodology; Writing – Review & Editing

Carolyn B. McNabb: Data Curation; Writing – Review & Editing

Derek K. Jones: Conceptualization; Methodology; Writing – Review & Editing; Supervision; Funding Acquisition

Andrew D. Lawrence: Conceptualization; Methodology; Writing – Review & Editing; Visualization

## Funding Information

This research was funded by a Wellcome Strategic Award [Grant number 104943/Z/14/Z].

Carl J. Hodgetts was supported by the Biotechnology and Biological Sciences Research Council (BBSRC) [Grant number BB/V010549/1].

Andrew D. Lawrence was supported by the Biotechnology and Biological Sciences Research Council (BBSRC) [Grant number BB/V008242/2].

## Conflicts of Interest

The authors report there are no competing interests to declare.

## Technical Terms

Myelin water fraction: MRI-derived estimate of water trapped within myelin sheaths.

Tractography: A computational method that uses diffusion-MRI data to reconstruct and visualize the white matter tracts of the brain.

Graph theory: Mathematical framework for analysing network organisation.

Characteristic path length: Average shortest distance between all pairs of network nodes.

Global efficiency: Measure of how efficiently information is exchanged across a network.

Mean clustering coefficient: Measure of local interconnectedness among neighbouring nodes.

Mean nodal strength: Average sum of the weights of the edges connected to the node.

## Notes

https://www.nature.com/articles/s41597-024-04154-7#Ack1

### Competing Interest Statement

The authors have declared no competing interest.

## References

Abraham, A., Schubotz, R. I., & von Cramon, D. Y. (2008). Thinking about the future versus the past in personal and non-personal contexts. Brain Research, 1233, 106–119. 10.1016/j.brainres.2008.07.084

Addis, D. R., & Schacter, D. L. (2008). Constructive episodic simulation: Temporal distance and detail of past and future events modulate hippocampal engagement. Hippocampus, 18(2), 227–237. 10.1002/hipo.20405

Addis, D. R., Valenzano, S. L., & Kinley, I. (2025). On future thinking: Specificity, reasoning, and schemas. Journal of Applied Research in Memory and Cognition, 14(1), 30–36. 10.1037/mac0000221

Addis, D. R., Wong, A. T., & Schacter, D. L. (2007). Remembering the past and imagining the future: Common and distinct neural substrates during event construction and elaboration. Neuropsychologia, 45(7), 1363–1377. 10.1016/j.neuropsychologia.2006.10.016

Addis, D. R., Wong, A. T., & Schacter, D. L. (2008). Age-related changes in the episodic simulation of future events. Psychological Science, 19(1), 33–41. 10.1111/j.1467-9280.2008.02043.x.

Ahmad, S., & Yap, P. T. (2026). Multiscale brain development across the human lifespan. Communications Biology, 9(1), 965. 10.1038/s42003-026-10565-6

Alexander-Bloch, A., Giedd, J. N., & Bullmore, E. (2013). Imaging structural co-variance between human brain regions. Nature Reviews Neuroscience, 14(5), 322–336. 10.1038/nrn3465

Andersson, J. L. R., & Sotiropoulos, S. N. (2016). An integrated approach to correction for off-resonance effects and subject movement in diffusion MR imaging. Neuroimage, 125, 1063–1078. 10.1016/j.neuroimage.2015.10.019

Andersson, J. L. R., Skare, S., & Ashburner, J. (2003). How to correct susceptibility distortions in spinecho echo-planar images: application to diffusion tensor imaging. Neuroimage, 20(2), 870–888. 10.1016/S1053-8119(03)00336-7

Andrews-Hanna, J. R., Reidler, J. S., Sepulcre, J., Poulin, R., & Buckner, R. L. (2010). Functional-anatomic fractionation of the brain’s default network. Neuron, 65(4), 550–562. 10.1016/j.neuron.2010.02.005

Andrews-Hanna, J. R. (2012). The Brain’s Default Network and Its Adaptive Role in Internal Mentation. The Neuroscientist, 18(3), 251–270. 10.1177/1073858411403316

Andrews-Hanna, J. R., Smallwood, J., & Spreng, R. N. (2014). The default network and self-generated thought: Component processes, dynamic control, and clinical relevance. Annals of the New York Academy of Sciences, 1316(1), 29–52. 10.1111/nyas.12360

Avants, B. B., Tustison, N., & Song, G. (2009). Advanced normalization tools (ANTS). The Insight Journal, 2(365), 1–35.

Bae, J., Joynes, C. M., Gong, Z., Duggan, M. R., Walker, K. A., & Bouhrara, M. (2026). Multi-omics validation of myelin water fraction as a myelin-specific MRI biomarker. Neuroimage, 330, 121852. 10.1016/j.neuroimage.2026.121852

Bar, M., & Aminoff, E. (2003). Cortical analysis of visual context. Neuron, 38(2), 347–358. 10.1016/S0896-6273(03)00167-3

Beaty, R. E., Kaufman, S. B., Benedek, M., Jung, R. E., Kenett, Y. N., Jauk, E., … & Silvia, P. J. (2016). Personality and complex brain networks: The role of openness to experience in default network efficiency. Human Brain Mapping, 37(2), 773–779. 10.1002/hbm.23065

Beaty, R. E., & Schacter, D. L. (2018). Episodic memory and cognitive control: Contributions to creative idea production. In R. E. Jung & O. Vartanian (Eds.), The Cambridge handbook of the neuroscience of creativity (pp. 249–260). Cambridge University Press. 10.1017/9781316556238.015

Bellana, B., Liu, Z.-X., Diamond, N. B., Grady, C. L., & Moscovitch, M. (2017). Similarities and differences in the default mode network across rest, retrieval, and future imagining. Human Brain Mapping, 38(3), 1155–1171. 10.1002/hbm.23445

Bells, S., Lefebvre, J., Prescott, S. A., Dockstader, C., Bouffet, E., Skocic, J., … & Mabbott, D. J. (2017). Changes in white matter microstructure impact cognition by disrupting the ability of neural assemblies to synchronize. The Journal of Neuroscience, 37(34), 8227–8238. 10.1523/JNEUROSCI.0481-17.2017

Benoit, R. G., & Schacter, D. L. (2015). Specifying the core network supporting episodic simulation and episodic memory by activation likelihood estimation. Neuropsychologia, 75, 450–457. 10.1016/j.neuropsychologia.2015.06.034

Berntsen, D., Hoyle, R. H., & Rubin, D. C. (2019). The autobiographical recollection test (ART): A measure of individual differences in autobiographical memory. Journal of Applied Research in Memory and Cognition, 8(3), 305–318. 10.1016/j.jarmac.2019.06.005

Braga R. M. (2025). The canonical default network comprises parallel distributed networks with distinct medial temporal lobe connections. Current Opinion in Behavioral Sciences, 66, 101610. 10.1016/j.cobeha.2025.101610

Bullmore, E., & Sporns, O. (2009). Complex brain networks: Graph theoretical analysis of structural and functional systems. Nature Reviews Neuroscience, 10(3), 186–198. 10.1038/nrn2575

Bullmore, E., & Sporns, O. (2012). The economy of brain network organization. Nature Reviews Neuroscience, 13(5), 336–349. 10.1038/nrn3214

Buyanova, I. S., & Arsalidou, M. (2021). Cerebral white matter myelination and relations to age, gender, and cognition: A selective review. Frontiers in Human Neuroscience, 15, 662031. 10.3389/fnhum.2021.662031

Wood, T. C. (2018). QUIT: quantitative imaging tools. The Journal of Open Source Software, 3(26), 656. 10.21105/joss.00656

Cai, Q., Thomas, H., Hyde, V., Laguna, P. L., McNabb, C. B., Singh, K. D., Jones, D. K., & Messaritaki, E. (2025). Decoding brain structure-function dynamics in health and in psychosis via an autoencoder. Scientific Reports, 15(1), 40052. 10.1038/s41598-025-24232-z

Cavanna, A. E., & Trimble, M. R. (2006). The precuneus: A review of its functional anatomy and behavioural correlates. Brain, 129(3), 564–583. 10.1093/brain/awl004.

Clark, I. A., & Maguire, E. A. (2020). Do questionnaires reflect their purported cognitive functions? Cognition, 195, 104114. 10.1016/j.cognition.2019.104114

Clark, I. A., Mohammadi, S., Callaghan, M. F., & Maguire, E. A. (2022). Conduction velocity along a key white matter tract is associated with autobiographical memory recall ability. ELife, 11. 10.7554/eLife.79303

Conti, F., & Irish, M. (2021). Harnessing Visual Imagery and Oculomotor Behaviour to Understand Prospection. Trends in Cognitive Sciences, 25(4), 272–283. 10.1016/j.tics.2021.01.009

Crawford, B., Muhlert, N., MacDonald, G., & Lawrence, A. D. (2020). Brain structure correlates of expected social threat and reward. Scientific Reports, 10(1), 18010. 10.1038/s41598-020-74334-z

Dale, A. M., & Sereno, M. I. (1993). Improved Localizadon of Cortical Activity by Combining EEG and MEG with MRI Cortical Surface Reconstruction: A Linear Approach. Journal of Cognitive Neuroscience, 5(2), 162–176. 10.1162/jocn.1993.5.2.162

Dale, A. M., Fischl, B., & Sereno, M. I. (1999). Cortical surface-based analysis. I. Segmentation and surface reconstruction. Neuroimage, 9(2), 179–194. 10.1006/nimg.1998.0395

Daviddi, S., Pedale, T., St Jacques, P. L., Schacter, D. L., & Santangelo, V. (2023). Common and distinct correlates of construction and elaboration of episodic-autobiographical memory: An ALE meta-analysis. Cortex, 163, 123–138. 10.1016/j.cortex.2023.03.005

Daviddi, S., Yaya, G., Sperduti, M., et al. (2025). A systematic review and meta-analysis of the neural correlates of direct vs. generative retrieval of episodic autobiographical memory. Neuropsychology Review, 35, 710–730. 10.1007/s11065-024-09653-3

Dawes, A. J., Keogh, R., Andrillon, T., & Pearson, J. (2020). A cognitive profile of multi sensory imagery, memory and dreaming in aphantasia. Scientific Reports, 10(1), 1–10. 10.1038/s41598-020-65705-7

Dawes, A. J., Keogh, R., Robuck, S., & Pearson, J. (2022). Memories with a blind mind: Remembering the past and imagining the future with aphantasia. Cognition, 227, 105192. 10.1016/j.cognition.2022.105192

Deoni, S. C. L. (2007). High-resolution T1 mapping of the brain at 3T with driven equilibrium single pulse observation of T1 with high-speed incorporation of RF field inhomogeneities (DESPOT1-HIFI). Journal of Magnetic Resonance Imaging, 26(4), 1106–1111. 10.1002/jmri.21130

Deoni, S. C. L., Matthews, L., & Kolind, S. H. (2013). One component? Two components? Three? The effect of including a nonexchanging “free” water component in multicomponent driven equilibrium single pulse observation of T1 and T2. Magnetic Resonance in Medicine, 70(1), 147–154. 10.1002/mrm.24429

Deoni, S. C. L., Rutt, B. K., & Peters, T. M. (2003). Rapid combined T1 and T2 mapping using gradient recalled acquisition in the steady state. Magnetic Resonance in Medicine, 49(3), 515–526. 10.1002/mrm.10407

Deoni, S. C. L., Rutt, B. K., Arun, T., Pierpaoli, C., & Jones, D. K. (2008). Gleaning multicomponent T1 and T2 information from steady-state imaging data. Magnetic Resonance in Medicine, 60(6), 1372–1387. 10.1002/mrm.21704

Desikan, R. S., Ségonne, F., Fischl, B., Quinn, B. T., Dickerson, B. C., Blacker, D., Buckner, R. L., Dale, A. M., Maguire, R. P., Hyman, B. T., Albert, M. S., & Killiany, R. J. (2006). An automated labeling system for subdividing the human cerebral cortex on MRI scans into gyral based regions of interest. Neuroimage, 31(3), 968–980. 10.1016/j.neuroimage.2006.01.021

Dhollander, T., Mito, R., Raffelt, D., & Connelly, A. (2019). Improved white matter response function estimation for 3-tissue constrained spherical deconvolution. International Society for Magnetic Resonance in Medicine (Vol. 27, p. 555).

Dhollander, T., Raffelt, D., & Connelly, A. (2016). Unsupervised 3-tissue response function estimation from single-shell or multi-shell diffusion MR data without a co-registered T1 image. ISMRM Workshop on Breaking the Barriers of Diffusion MRI, 5.

Dikmen, S. S., Bauer, P. J., Weintraub, S., Mungas, D., Slotkin, J., Beaumont, J. L., Gershon, R., Temkin, N. R., & Heaton, R. K. (2014). Measuring episodic memory across the lifespan: NIH Toolbox Picture Sequence Memory Test. Journal of the International Neuropsychological Society, 20(6), 611–619. 10.1017/S1355617714000460

Drakesmith, M., Harms, R., Rudrapatna, S. U., Parker, G. D., Evans, C. J., & Jones, D. K. (2019). Estimating axon conduction velocity in vivo from microstructural MRI. Neuroimage, 203, 116186. 10.1016/j.neuroimage.2019.116186

Edmonds, D., Salvo, J. J., Anderson, N., Lakshman, M., Yang, Q., Kay, K., … & Braga, R. M. (2024). The human social cognitive network contains multiple regions within the amygdala. Science Advances, 10(47), eadp0453. 10.1126/sciadv.adp0453

Fan, C. L., Abdi, H., & Levine, B. (2021). On the relationship between trait autobiographical episodic memory and spatial navigation. Memory and Cognition, 49(2), 265–275. 10.3758/s13421-020-01093-7

Faul, F., Erdfelder, E., Lang, AG. et al. G*Power 3: A flexible statistical power analysis program for the social, behavioral, and biomedical sciences. Behavior Research Methods 39, 175–191 (2007). 10.3758/BF03193146

Fischl, B., & Dale, A. M. (2000). Measuring the thickness of the human cerebral cortex from magnetic resonance images. Proceedings of the National Academy of Sciences of the United States of America, 97(20), 11050–11055. 10.1073/pnas.200033797

Fischl, B., Liu, A., & Dale, A. M. (2001). Automated manifold surgery: constructing geometrically accurate and topologically correct models of the human cerebral cortex. IEEE Transactions on Medical Imaging, 20(1), 70–80. 10.1109/42.906426

Fischl, B., Salat, D. H., van der Kouwe, A. J. W., Makris, N., Ségonne, F., Quinn, B. T., & Dale, A. M. (2004). Sequence-independent segmentation of magnetic resonance images. Neuroimage, 23 Suppl 1, S69–84. 10.1016/j.neuroimage.2004.07.016

Fischl, B., Sereno, M. I., & Dale, A. M. (1999). Cortical surface-based analysis. II: Inflation, flattening, and a surface-based coordinate system. Neuroimage, 9(2), 195–207. 10.1006/nimg.1998.0396

Fischl, B., Sereno, M. I., Tootell, R. B., & Dale, A. M. (1999). High-resolution intersubject averaging and a coordinate system for the cortical surface. Human Brain Mapping, 8(4), 272–284. 10.1002/(sici)1097-0193(1999)8:4<272::aid-hbm10>3.0.co;2-4

Fischl, B., Salat, D. H., Busa, E., Albert, M., Dieterich, M., Haselgrove, C., van der Kouwe, A., Killiany, R., Kennedy, D., Klaveness, S., Montillo, A., Makris, N., Rosen, B., & Dale, A. M. (2002). Whole brain segmentation: automated labeling of neuroanatomical structures in the human brain. Neuron, 33(3), 341–355. 10.1016/s0896-6273(02)00569-x

Fivush, R. (2011). The development of autobiographical memory. Annual Review of Psychology, 62, 559–582. 10.1146/annurev.psych.121208.131702

Geib, B. R., Stanley, M. L., Wing, E. A., Laurienti, P. J., & Cabeza, R. (2017). Hippocampal contributions to the large-scale episodic memory network predict vivid visual memories. Cerebral Cortex, 27(1), 680–693. 10.1093/cercor/bhv272

Geib, B.R., Stanley, M.L., Dennis, N.A., Woldorff, M.G. and Cabeza, R. (2017), From hippocampus to whole-brain: The role of integrative processing in episodic memory retrieval. Human Brain Mapping, 38, 2242–2259. 10.1002/hbm.23518

Gignac, G. E., & Szodorai, E. T. (2016). Effect size guidelines for individual differences researchers. Personality and individual differences, 102, 74–78. 10.1016/j.paid.2016.06.069

Gilboa, A., & Marlatte, H. (2017). Neurobiology of Schemas and Schema-Mediated Memory. Trends in Cognitive Sciences, 21(8), 618–631. 10.1016/j.tics.2017.04.013

Gilmore, A. W., Nelson, S. M., Chen, H. Y., & McDermott, K. B. (2018). Task-related and resting-state fMRI identify distinct networks that preferentially support remembering the past and imagining the future. Neuropsychologia, 110, 180–189. 10.1016/j.neuropsycholo-gia.2017.06.016

Girn, M., Setton, R., Turner, G. R., & Spreng, R. N. (2024). The “limbic network,” comprising orbitofrontal and anterior temporal cortex, is part of an extended default network: Evidence from multi-echo fMRI. Network Neuroscience, 8(3), 860–882. 10.1162/netn_a_00385

Gur, R. C., Richard, J., Hughett, P., Calkins, M. E., Macy, L., Bilker, W. B., Brensinger, C., & Gur, R. E. (2010). A cognitive neuroscience-based computerized battery for efficient measurement of individual differences: standardization and initial construct validation. Journal of Neuroscience Methods, 187(2), 254–262. 10.1016/j.jneumeth.2009.11.017

Habermas, T., & Diel, V. (2013). The episodicity of verbal reports of personally significant autobiographical memories: Vividness correlates with narrative text quality more than with detailedness or memory specificity. Frontiers in Behavioral Neuroscience, 7, 110. 10.3389/fnbeh.2013.00110

Hadi, A. S., Imon, A. H. M. R., & Werner, M. (2009). Detection of outliers. Wiley Interdisciplinary Reviews: Computational Statistics, 1(1), 57–70. 10.1002/wics.6

Han, X., Jovicich, J., Salat, D., van der Kouwe, A., Quinn, B., Czanner, S., Busa, E., Pacheco, J., Albert, M., Killiany, R., Maguire, P., Rosas, D., Makris, N., Dale, A., Dickerson, B., & Fischl, B. (2006). Reliability of MRI-derived measurements of human cerebral cortical thickness: the effects of field strength, scanner upgrade and manufacturer. Neuroimage, 32(1), 180–194. 10.1016/j.neuroimage.2006.02.051

Hassabis, D., & Maguire, E. A. (2007). Deconstructing episodic memory with construction. Trends in Cognitive Sciences, 11(7), 299–306. 10.1016/j.tics.2007.05.001

Hassabis, D., Kumaran, D., Vann, S. D., & Maguire, E. A. (2007). Patients with hippocampal amnesia cannot imagine new experiences. Proceedings of the National Academy of Sciences of the United States of America, 104(5), 1726–1731. 10.1073/pnas.0610561104

Hodgetts, C. J., Postans, M., Warne, N., Varnava, A., Lawrence, A. D., & Graham, K. S. (2017). Distinct contributions of the fornix and inferior longitudinal fasciculus to episodic and semantic autobiographical memory. Cortex, 94, 1–14. 10.1016/j.cortex.2017.05.010

Irish, M., & Piguet, O. (2013). The pivotal role of semantic memory in remembering the past and imagining the future. Frontiers in Behavioral Neuroscience, 7, 27. 10.3389/fnbeh.2013.00027

Irish, M., Addis, D. R., Hodges, J. R., & Piguet, O. (2012a). Considering the role of semantic memory in episodic future thinking: evidence from semantic dementia. Brain: A Journal of Neurology, 135(Pt 7), 2178–2191. 10.1093/brain/aws119

Irish, M., Addis, D. R., Hodges, J. R., & Piguet, O. (2012b). Exploring the content and quality of episodic future simulations in semantic dementia. Neuropsychologia, 50(14), 3488–3495. 10.1016/j.neuropsychologia.2012.09.014

Isensee, F., Schell, M., Pflueger, I., Brugnara, G., Bonekamp, D., Neuberger, U., Wick, A., Schlemmer, H.-P., Heiland, S., Wick, W., Bendszus, M., Maier-Hein, K. H., & Kickingereder, P. (2019). Automated brain extraction of multisequence MRI using artificial neural networks. Human Brain Mapping, 40(17), 4952–4964. 10.1002/hbm.24750

Jenkinson, M., & Smith, M. A. (2001). A global optimisation method for affine registration of brain images. Medical Image Analysis, 5(2), 143–156. 10.1016/s1361-8415(01)00036-6

Jenkinson, M., Bannister, P. R., Brady, J. M., & Smith, S. M. (2002). Improved optimisation for the robust and accurate linear registration and motion correction of brain images. Neuroimage, 17(2), 825–841. 10.1016/s1053-8119(02)91132-8

Jenkinson, M., Beckmann, C. F., Behrens, T. E., Woolrich, M. W., & Smith, S. M. (2012). FSL. Neuroimage, 62(2), 782–790. 10.1016/j.neuroimage.2011.09.015

Jeurissen, B., Tournier, J. D., Dhollander, T., Connelly, A., & Sijbers, J. (2014). Multi-tissue constrained spehrical deconvolution for improved analysis of multi-shell diffusion MRI data. Neuroimage, 103, 411–426. 10.1016/j.neuroimage.2014.07.061

Jones, D. K., Horsfield, M. A., & Simmons, A. (1999). Optimal strategies for measuring diffusion in anisotropic systems by magnetic resonance imaging. Magnetic Resonance in Medicine, 42(3), 515–525. 10.1002/(SICI)1522-2594(199909)42:3<515::AID-MRM14>3.0.CO;2-Q

Jones, D. K., Knösche, T. R., & Turner, R. (2013). White matter integrity, fiber count, and other fallacies: the do’s and don’ts of diffusion MRI. Neuroimage, 73, 239–254. 10.1016/j.neuroimage.2012.06.081

Jovicich, J., Czanner, S., Greve, D., Haley, E., van der Kouwe, A., Gollub, R., Kennedy, D., Schmitt, F., Brown, G., Macfall, J., Fischl, B., & Dale, A. (2006). Reliability in multi-site structural MRI studies: effects of gradient non-linearity correction on phantom and human data. Neuroimage, 30(2), 436–443. 10.1016/j.neuroimage.2005.09.046

Jung, J., Cloutman, L. L., Binney, R. J., & Ralph, M. A. L. (2017). The structural connectivity of higher order association cortices reflects human functional brain networks. Cortex, 97, 221–239. 10.1016/j.cortex.2016.08.011

Karahan, E., Tait, L., Si, R., Özkan, A., Szul, M. J., Graham, K. S., Lawrence, A. D., & Zhang, J. (2022). The interindividual variability of multimodal brain connectivity maintains spatial heterogeneity and relates to tissue microstructure. Communications Biology, 5(1), 1007. 10.1038/s42003-022-03974-w

Kellner, E., Dhital, B., Kiselev, V. G., & Reisert, M. (2016). Gibbs-ringing artifact removal based on local subvoxel-shifts. Magnetic Resonance in Medicine, 76(5), 1574–1581. 10.1002/mrm.26054

Kensinger, E. A., & Fields, E. (2024). Affective memory. In Oxford handbook of human memory: Foundations and applications (pp. 553–584). Oxford University Press.

Kensinger, E. A., Addis, D. R., & Atapattu, R. K. (2011). Amygdala activity at encoding corresponds with memory vividness and with memory for select episodic details. Neuropsychologia, 49(4), 663–673. 10.1016/j.neuropsychologia.2011.01.017

Kerns, J. G., Cohen, J. D., MacDonald, A. W., III, Cho, R. Y., Stenger, V. A., & Carter, C. S. (2004). Anterior cingulate conflict monitoring and adjustments in control. Science, 303(5660), 1023–1026. 10.1126/science.1089910.

Kucyi, A., & Davis, K. D. (2014). Dynamic functional connectivity of the default mode network tracks day-dreaming. Neuroimage, 100, 471–480. 10.1016/j.neuroimage.2014.06.044

Lechowicz, M., Miller, L., Irish, M., Addis, D. R., Mohamed, A., & Lah, S. (2016). Imagining future events in patients with unilateral temporal lobe epilepsy. The British Journal of Clinical Psychology, 55(2), 187–205. 10.1111/bjc.12107

LePort, A. K. R., Mattfeld, A. T., Dickinson-Anson, H., Fallon, J. H., Stark, C. E. L., Kruggel, F., Cahill, L., & McGaugh, J. L. (2012). Behavioral and neuroanatomical investigation of Highly Superior Autobiographical Memory (HSAM). Neurobiology of Learning and Memory, 98(1), 78–92. 10.1016/j.nlm.2012.05.002

Levine, B., Svoboda, E., Hay, J. F., Winocur, G., & Moscovitch, M. (2002). Aging and autobiographical memory: dissociating episodic from semantic retrieval. Psychology and Aging, 17(4), 677–689. 10.1037//0882-7974.17.4.677

Li, J., Curley, W. H., Guerin, B., Dougherty, D. D., Dalca, A. V., Fischl, B., Horn, A., & Edlow, B. L. (2021). Mapping the subcortical connectivity of the human default mode network. Neuroimage, 245, 118758. 10.1016/j.neuroimage.2021.118758

Matyi, M. A., & Spielberg, J. M. (2022). The structural brain network topology of episodic memory. Plos One, 17(6), e0270592. 10.1371/journal.pone.0270592

McDermott, K. B., Szpunar, K. K., & Christ, S. E. (2009). Laboratory-based and autobiographical retrieval tasks differ substantially in their neural substrates. Neuropsychologia, 47(11), 2290–2298. 10.1016/j.neuropsychologia.2008.12.025

McLelland, V. C., Devitt, A. L., Schacter, D. L., & Addis, D. R. (2015). The past, the present, and the future: a meta-analysis of functional neuroimaging data associated with episodic memory and simulation. Journal of Cognitive Neuroscience, 27(7), 1432–1447. 10.1162/JOCN_a_00810

McNabb, C. B., Driver, I. D., Hyde, V., Hughes, G., Chandler, H. L., Thomas, H., Allen, C., Messaritaki, E., Hodgetts, C. J., Hedge, C., Engel, M., Standen, S. F., Morgan, E. L., Stylianopoulou, E., Manolova, S., Reed, L., Ploszajski, M., Drakesmith, M., Germuska, M., … Jones, D. K. (2025). WAND: A multi-modal dataset integrating advanced MRI, MEG, and TMS for multi-scale brain analysis. Scientific Data, 12(1), 220. 10.1038/s41597-024-04154-7

Menon, V. (2023). 20 years of the default mode network: A review and synthesis. Neuron, 111(16), 2469–2487. 10.1016/j.neuron.2023.04.023

Messaritaki, E., Foley, S., Schiavi, S., Magazzini, L., Routley, B., Jones, D. K., & Singh, K. D. (2021). Predicting MEG resting-state functional connectivity from microstructural information. Network Neuroscience, 5(2), 477–504. 10.1162/netn_a_00187

Messaritaki, E., Hedge, C., Laguna, P. L., McNabb, C. B., Jones, D., & Sumner, P. (2025). Composite reaction time, not evidence accumulation efficiency, correlates with whole-brain white matter characteristics. Imaging Neuroscience 2026; 4, IMAG.a.1303. 10.1162/IMAG.a.1303

Messaritaki, E., Hedge, C., Sumner, P., & Jones, D. (2023). Cognitive analysis of the Welsh Advanced Neuroimaging Database (WAND). Open Science Framework. 10.17605/OSF.IO/SNYQF

Mesulam, M. M. (1990). Large-scale neurocognitive networks and distributed processing for attention, language, and memory. Annals of Neurology, 28(5), 597–613. 10.1002/ana.410280502.

Mišić, B., Sporns, O., & McIntosh, A. R. (2014). Communication efficiency and congestion of signal traffic in large-scale brain networks. PLoS Computational Biology, 10(1), e1003427. 10.1371/journal.pcbi.1003427

Murray, E. A., Moylan, E. J., Saleem, K. S., Basile, B. M., & Turchi, J. (2015). Specialized areas for value updating and goal selection in the primate orbitofrontal cortex. eLife, 4, e11695. 10.7554/eLife.11695.

Nelson, M. C., Da Lu, W., Leppert, I. R., Hansen, H. A., Rowley, C. D., Misic, B., & Tardif, C. L. (2026). The role of white matter myelin in structural-functional network coupling. Communications Biology 9, 488 (2026). 10.1038/s42003-026-09813-6

Nunnally, J. C. (1978). Psychometric theory (2nd ed.). McGraw-Hill.

Østby, Y., Walhovd, K. B., Tamnes, C. K., Grydeland, H., Westlye, L. T., & Fjell, A. M. (2012). Mental time travel and default-mode network functional connectivity in the developing brain. Proceedings of the National Academy of Sciences of the United States of America, 109(42), 16800–16804. 10.1073/pnas.1210627109

Owen, J. P., Ziv, E., Bukshpun, P., Pojman, N., Wakahiro, M., Berman, J. I., Roberts, T. P., Friedman, E. J., Sherr, E. H., & Mukherjee, P. (2013). Test-retest reliability of computational network measurements derived from the structural connectome of the human brain. Brain Connectivity, 3(2), 160–176. 10.1089/brain.2012.0121

Palombo, D. J., Alain, C., Söderlund, H., Khuu, W., & Levine, B. (2015). Severely deficient autobiographical memory (SDAM) in healthy adults: A new mnemonic syndrome. Neuropsychologia, 72, 105–118. 10.1016/j.neuropsychologia.2015.04.012

Palombo, D. J., Sheldon, S., & Levine, B. (2018). Individual differences in autobiographical memory. Trends in Cognitive Sciences, 22(7), 583–597. 10.1016/j.tics.2018.04.007

Palombo, D. J., Williams, L. J., Abdi, H., & Levine, B. (2013). The survey of autobiographical memory (SAM): a novel measure of trait mnemonics in everyday life. Cortex, 49(6), 1526–1540. 10.1016/j.cortex.2012.08.023

Petersen, S. E., & Sporns, O. (2015). Brain Networks and Cognitive Architectures. Neuron, 88(1), 207–219. 10.1016/j.neuron.2015.09.027

Petrican, R., Palombo, D. J., Sheldon, S., & Levine, B. (2020). The neural dynamics of individual differences in episodic autobiographical memory. ENeuro, 7(2). 10.1523/ENEURO.0531-19.2020

Phelps, E. A., & LeDoux, J. E. (2005). Contributions of the amygdala to emotion processing: From animal models to human behavior. Neuron, 48(2), 175–187. 10.1016/j.neu-ron.2005.09.025

Phelps, E. A., & Sharot, T. (2008). How (and Why) Emotion Enhances the Subjective Sense of Recollection. Current Directions in Psychological Science, 17(2), 147–152. 10.1111/j.1467-8721.2008.00565.x

Picco, S., Pedreira, M. E., & Fernández, R. S. (2020). Psychometric validation of the Survey of Autobiographical Memory: confirmatory factor analysis and network analysis. Memory, 28(8), 1037–1050. 10.1080/09658211.2020.1812662

Pierpaoli, C., Jezzard, P., Basser, P. J., Barnett, A., & Di Chiro, G. (1996). Diffusion tensor MR imaging of the human brain. Radiology, 201(3), 637–648. 10.1148/radiology.201.3.8939209

Preminger, S., Harmelech, T., & Malach, R. (2011). Stimulus-free thoughts induce differential activation in the human default network. Neuroimage, 54(2), 1692–1702. 10.1016/j.neuroimage.2010.09.055

Ray, K.L., Ragland, J.D., MacDonald, A.W. et al. Dynamic reorganization of the frontal parietal network during cognitive control and episodic memory. Cognitive, Affective, & Behavioral Neuroscience, 20, 76–90 (2020). 10.3758/s13415-019-00753-9

Rayner, G., Antoniou, M., Jackson, G., & Tailby, C. (2022). Compromised future thinking: another cognitive cost of temporal lobe epilepsy. Brain Communications, 4(2), fcac062. 10.1093/braincomms/fcac062

Reisberg, D., Heuer, F., McLean, J., & O’Shaughnessy, M. (1988). The quantity, not the quality, of affect predicts memory vividness. Bulletin of the Psychonomic Society, 26(2), 100–103. 10.3758/BF03334873

Renoult, L., Irish, M., Moscovitch, M., & Rugg, M. D. (2019). From Knowing to Remembering: The Semantic-Episodic Distinction. Trends in Cognitive Sciences, 23(12), 1041–1057. 10.1016/j.tics.2019.09.008

Reuter, M., Rosas, H. D., & Fischl, B. (2010). Highly accurate inverse consistent registration: a robust approach. Neuroimage, 53(4), 1181–1196. 10.1016/j.neuroimage.2010.07.020

Reuter, M., Schmansky, N. J., Rosas, H. D., & Fischl, B. (2012). Within-subject template estimation for unbiased longitudinal image analysis. Neuroimage, 61(4), 1402–1418. 10.1016/j.neuroimage.2012.02.084

Roediger III, H. L., & McDermott, K. B. (2013). Two types of event memory. Proceedings of the National Academy of Sciences, 110(52), 20856–20857. 10.1073/pnas.1321373110

Rubin, D. C. (2021). Properties of autobiographical memories are reliable and stable individual differences. Cognition, 210, 104583. 10.1016/j.cognition.2021.104583

Rubinov, M., & Sporns, O. (2010). Complex network measures of brain connectivity: uses and interpretations. Neuroimage, 52(3), 1059–1069. 10.1016/j.neuroimage.2009.10.003

Rugg, M. D., & Vilberg, K. L. (2013). Brain networks underlying episodic memory retrieval. Current Opinion in Neurobiology, 23(2), 255–260. 10.1016/j.conb.2012.11.005

Sairanen, V., Leemans, A., & Tax, C. M. W. (2018). Fast and accurate Slicewise OutLIer Detection (SOLID) with informed model estimation for diffusion MRI data. Neuroimage, 181, 331–346. 10.1016/j.neuroimage.2018.07.046

Schacter, D. L., & Addis, D. R. (2007). On the constructive episodic simulation of past and future events. Behavioral and Brain Sciences, 30(3), 331–332. 10.1017/S0140525X07002178

Schacter, D. L., & Thakral, P. P. (2024). Constructive Memory and Conscious Experience. Journal of Cognitive Neuroscience, 36(8), 1567–1577. 10.1162/jocn_a_02201

Schacter, D. L., Addis, D. R., Hassabis, D., Martin, V. C., Spreng, R. N., & Szpunar, K. K. (2012). The future of memory: remembering, imagining, and the brain. Neuron, 76(4), 677–694. 10.1016/j.neuron.2012.11.001

Schacter, D. L., Benoit, R. G., & Szpunar, K. K. (2017). Episodic future thinking: mechanisms and functions. Current Opinion in Behavioral Sciences, 17, 41–50. 10.1016/j.cobeha.2017.06.013

Schedlbauer, A., Copara, M., Watrous, A. et al. Multiple interacting brain areas underlie successful spatiotemporal memory retrieval in humans. Scientific Reports, 4, 6431 (2014). 10.1038/srep06431

Ségonne, F., Dale, A. M., Busa, E., Glessner, M., Salat, D., Hahn, H. K., & Fischl, B. (2004). A hybrid approach to the skull stripping problem in MRI. Neuroimage, 22(3), 1060–1075. 10.1016/j.neuroimage.2004.03.032

Sergerie, K., Chochol, C., & Armony, J. L. (2008). The role of the amygdala in emotional processing: A quantitative meta-analysis of functional neuroimaging studies. Neuroscience & Biobehavioral Reviews, 32(4), 811–830. 10.1016/j.neubiorev.2007.12.002

Setton, R., Lockrow, A. W., Turner, G. R., & Spreng, R. N. (2022). Troubled past: A critical psychometric assessment of the self-report Survey of Autobiographical Memory (SAM). Behavior Research Methods, 54(1), 261–286. 10.3758/s13428-021-01604-7

Shapira-Lichter, I., Oren, N., Jacob, Y., Gruberger, M., & Hendler, T. (2013). Portraying the unique contribution of the default mode network to internally driven mnemonic processes. Proceedings of the National Academy of Sciences, 110(13), 4950–4955. 10.1073/pnas.1209888110

Sheldon, S., Farb, N., Palombo, D. J., & Levine, B. (2016). Intrinsic medial temporal lobe connectivity relates to individual differences in episodic autobiographical remembering. Cortex, 74, 206–216. 10.1016/j.cortex.2015.11.005

Smith, R. E., Tournier, J.-D., Calamante, F., & Connelly, A. (2012). Anatomically-constrained tractography: improved diffusion MRI streamlines tractography through effective use of anatomical information. Neuroimage, 62(3), 1924–1938. 10.1016/j.neuroimage.2012.06.005

Smith, R. E., Tournier, J.-D., Calamante, F., & Connelly, A. (2015). SIFT2: Enabling dense quantitative assessment of brain white matter connectivity using streamlines tractography. Neuroimage, 119, 338–351. 10.1016/j.neuroimage.2015.06.092

Smith, S M. (2002). Fast robust automated brain extraction. Human Brain Mapping, 17(3), 143–155. 10.1002/hbm.10062

Smith, S. M., Jenkinson, M., Woolrich, M. W., Beckmann, C. F., Behrens, T. E. J., Johansen-Berg, H., Bannister, P. R., De Luca, M., Drobnjak, I., Flitney, D. E., Niazy, R. K., Saunders, J., Vickers, J., Zhang, Y., De Stefano, N., Brady, J. M., & Matthews, P. M. (2004). Advances in functional and structural MR image analysis and implementation as FSL. Neuroimage, 23(Suppl. 1), S208–S219. 10.1016/j.neuroimage.2004.07.051

Sporns, O. (2010). Networks of the brain. MIT Press, Cambridge.

Sporns, O. (2013). Network attributes for segregation and integration in the human brain. Current Opinion in Neurobiology, 23(2), 162–171. 10.1016/j.conb.2012.12.016

Spreng, R. N., Mar, R. A., & Kim, A. S. (2009). The common neural basis of autobiographical memory, prospection, navigation, theory of mind, and the default mode: a quantitative meta-analysis. Journal of Cognitive Neuroscience, 21(3), 489–510. 10.1162/jocn.2008.21029

Suárez, L. E., Markello, R. D., Betzel, R. F., & Misic, B. (2020). Linking Structure and Function in Macroscale Brain Networks. Trends in Cognitive Sciences, 24(4), 302–315. 10.1016/j.tics.2020.01.008

Suminaite, D., Lyons, D. A., & Livesey, M. R. (2019). Myelinated axon physiology and regulation of neural circuit function. Glia, 67(11), 2050–2062. 10.1002/glia.23603

Sylvester, C. M., Corbetta, M., Raichle, M. E., Rodebaugh, T. L., Schlaggar, B. L., Sheline, Y. I., Zorumski, C. F., & Lenze, E. J. (2012). Functional network dysfunction in anxiety and anxiety disorders. Trends in Neurosciences, 35(9), 527–535. 10.1016/j.tins.2012.04.012

Sylvester, C. M., Yu, Q., Srivastava, A. B., Marek, S., Zheng, A., Alexopoulos, D., Smyser, C. D., Shimony, J. S., Ortega, M., Dierker, D. L., Patel, G. H., Nelson, S. M., Gilmore, A. W., McDermott, K. B., Berg, J. J., Drysdale, A. T., Perino, M. T., Snyder, A. Z., Raut, R. V., Laumann, T. O., … Dosenbach, N. U. F. (2020). Individual-specific functional connectivity of the amygdala: A substrate for precision psychiatry. Proceedings of the National Academy of Sciences of the United States of America, 117(7), 3808–3818. 10.1073/pnas.1910842117 https://doi.org/10.1073/pnas.1910842117

Talarico, J. M., LaBar, K. S., & Rubin, D. C. (2004). Emotional intensity predicts autobiographical memory experience. Memory & Cognition, 32, 1118–1132. 10.3758/BF03196886

Talbot, J., Convertino, G., De Marco, M., Venneri, A., & Mazzoni, G. (2025). Highly superior autobiographical memory (HSAM): A systematic review. Neuropsychology Review, 35(1), 54–76. 10.1007/s11065-024-09632-8

Teghil, A., Bonavita, A., Procida, F., Giove, F., & Boccia, M. (2022). Temporal Organization of Episodic and Experience-near Semantic Autobiographical Memories: Neural Correlates and Context-dependent Connectivity. Journal of Cognitive Neuroscience, 34(12), 2256–2274. 10.1162/jocn_a_01906

Tournier, J.-D., Smith, R., Raffelt, D., Tabbara, R., Dhollander, T., Pietsch, M., Christiaens, D., Jeurissen, B., Yeh, C.-H., & Connelly, A. (2019). MRtrix3: A fast, flexible and open software framework for medical image processing and visualisation. Neuroimage, 202, 116137. 10.1016/j.neuroimage.2019.116137

Tulving E. (2001). Episodic memory and common sense: how far apart?. Philosophical Transactions of the Royal Society of London. Series B, Biological Sciences, 356(1413), 1505–1515. 10.1098/rstb.2001.0937

Tulving, E. (2002). Episodic memory: from mind to brain. Annual Review of Psychology, 53, 1–25. 10.1146/annurev.psych.53.100901.135114

Vann, S. D., Aggleton, J. P., & Maguire, E. A. (2009). What does the retrosplenial cortex do? Nature Reviews Neuroscience, 10(11), 792–802. 10.1038/nrn2733

Vatansever, D. (2026). Mental navigation as a default computational mode in human cognition. Current Opinion in Behavioral Sciences, 70, 101671. 10.1016/j.cobeha.2026.101671

Veraart, J., Fieremans, E., & Novikov, D. S. (2016). Diffusion MRI noise mapping using random matrix theory. Magnetic Resonance in Medicine, 76(5), 1582–1593. 10.1002/mrm.26059

Veraart, J., Novikov, D. S., Christiaens, D., Ades-Aron, B., Sijbers, J., & Fieremans, E. (2016). Denoising of diffusion MRI using random matrix theory. Neuroimage, 142, 394–406. 10.1016/j.neuroimage.2016.08.016

Villena-González, M., Wang, H.-T., Sormaz, M., Mollo, G., Margulies, D. S., Jefferies, E. A., & Smallwood, J. (2018). Individual variation in the propensity for prospective thought is associated with functional integration between visual and retrosplenial cortex. Cortex, 99, 224–234. 10.1016/j.cortex.2017.11.015

Visser, M., Jefferies, E., & Lambon Ralph, M. A. (2010). Semantic processing in the anterior temporal lobes: A meta-analysis of the functional neuroimaging literature. Journal of Cognitive Neuroscience, 22(6), 1083–1094. 10.1162/jocn.2009.21309

Weymar, M., & Schwabe, L. (2016). Amygdala and emotion: The bright side of it. Frontiers in Neuroscience, 10, 224. 10.3389/fnins.2016.00224

Williams, A. N., Ridgeway, S., Postans, M., Graham, K. S., Lawrence, A. D., & Hodgetts, C. J. (2020). The role of the pre-commissural fornix in episodic autobiographical memory and simulation. Neuropsychologia, 142, 107457. 10.1016/j.neuropsychologia.2020.107457

Xu, X., Yuan, H. & Lei, X. (2016). Activation and connectivity within the default mode network contribute independently to future-oriented thought. Scientific Reports, 6, 21001. 10.1038/srep21001

Yeh, C. H., Jones, D. K., Liang, X., Descoteaux, M., & Connelly, A. (2021). Mapping structural connectivity using diffusion MRI: challenges and opportunities. Journal of Magnetic Resonance Imaging, 53(6), 1666–1682. 10.1002/jmri.27188

Yeshurun, Y., Nguyen, M., & Hasson, U. (2021). The default mode network: where the idiosyncratic self meets the shared social world. Nature Reviews Neuroscience, 22(3), 181–192. 10.1038/s41583-020-00420-w

